# HP1B is a euchromatic Drosophila HP1 homolog with links to metabolism

**DOI:** 10.1101/372094

**Authors:** Benjamin B. Mills, Andrew D. Thomas, Nicole C. Riddle

## Abstract

Heterochromatin Protein 1 (HP1) proteins are an important family of chromosomal proteins conserved among all major eukaryotic lineages. While HP1 proteins are best known for their role in heterochromatin, many HP1 proteins function in euchromatin as well. As a group, HP1 proteins carry out diverse functions, playing roles in the regulation of gene expression, genome stability, chromatin structure, and DNA repair. While the heterochromatic HP1 proteins are well studied, our knowledge of HP1 proteins with euchromatic distribution is lagging behind. We have created the first mutations in HP1B, a Drosophila HP1 protein with euchromatic function, and the Drosophila homolog most closely related to mammalian HP1α, HP1β, and HP1γ. We find that HP1B is a non-essential protein in Drosophila, with mutations affecting fertility and animal activity levels. In addition, animals lacking HP1B show altered food intake and higher body fat levels. Gene expression analysis of animals lacking HP1B demonstrates that genes with functions in various metabolic processes are affected primarily by HP1B loss. Our findings suggest that there is a link between the chromatin protein HP1B and the regulation of metabolism.

## INTRODUCTION

Regulation of gene expression is an essential process. Errors in gene regulation lead to aberrant transcription or silencing, which in turn impact the fitness, health, and survival of the organism. In eukaryotes, chromatin, the composite of DNA and its associated proteins, mediates important aspects of gene regulation. Specific chromatin types are formed depending on the positioning of nucleosomes, the post-translational modification of the histones within the nucleosome, and the types of chromosomal proteins present. The simplest models distinguish two basic chromatin types, heterochromatin and euchromatin, but more elaborate models exist as well [1-3]. Generally speaking, euchromatin encompasses gene-rich, transcriptionally active sequences, while heterochromatin contains highly condensed, repeat-rich, transcriptionally inactive regions [4]. These chromatin types are biochemically distinct, and chromosomal proteins carry out a variety of functions in specifying the distinct chromatin structures. Some are histone modifying enzymes, nucleosome remodelers, or boundary proteins, while others serve as structural components of chromatin. Regardless of their specific molecular function, chromosomal proteins are crucial for gene regulation.

The Heterochromatin Protein 1 (HP1) proteins are a family of chromosomal proteins that carry out roles in gene regulation, genome stability, and DNA repair [5, 6]. HP1 proteins have been linked to cancer, and they also appear to contribute to cellular senescence [7-9]. The protein family is highly conserved, and HP1 proteins are found in the plant, animal, and fungal lineages (TFL2 in Arabidopsis, SWI6 in *S. pombe*, and HP1α, HP1β, and HP1γ in humans) [10, 11]. Two conserved domains characterize HP1 family proteins, an N-terminal chromo domain and a C-terminal chromo shadow domain [12]. A variable hinge domain connects these two conserved domains, and N- and/or C-terminal tails exist as well. The chromo domain is important for interactions of HP1 proteins with the chromatin (via its interaction with histone 3 methylated at lysine 9 [13-18]) and is critical for localization to specific genomic regions [6, 11]. Other parts of the HP1 proteins, however, appear to contribute to the genomic localization as well [19]. The chromo shadow domain can mediate dimerization between two HP1 molecules [20-22]. It is unclear however, if all HP1 proteins form dimers, be they homodimers or heterodimers. For HP1 molecules that dimerize, the protein surface created by dimerization is important for the interaction with other protein partners [21, 23]. Via the two conserved domains and various protein interactions, HP1 family members are able to contribute to many cellular processes, including transcription regulation.

Most eukaryotic genomes contain several genes encoding HP1 proteins. Usually, some of these proteins exhibit the characteristics that one typically associates with the term “heterochromatin protein” – a structural component of heterochromatin that has a silencing effect on gene expression [24, 25]. These members of the HP1 family tend to be encoded by essential genes, as the proteins are required for centromere function and loss leads to cell division defects [24]. If additional HP1 paralogs are present, they tend to have functions separate from (and not redundant with) the classical heterochromatin function.

Their localization pattern in the genome is distinct, as is their impact on gene regulation. For example, both humans and Drosophila have three somatic HP1 proteins [in addition, Drosophila has two germline-specific HP1s [26] and several partial HP1 derivatives [27]], HP1a, HP1B, and HP1C in Drosophila, and HP1α, HP1β, and HP1γ in humans (CBX5, CBX1, and CBX3). HP1a and HP1α are predominantly associated with heterochromatic regions of the genome [25, 28]. HP1B and HP1β are described as localizing to euchromatin and heterochromatin, while localization of HP1C and HP1γ is reported as predominantly euchromatic [19, 28]. While the role of HP1 proteins in heterochromatin formation is well understood, less is known about the function of HP1 proteins in euchromatin.

HP1B in Drosophila localizes to heterochromatin and euchromatin and can serve as model for the HP1 proteins that do not show the classical heterochromatic localization. It is also the Drosophila HP1 paralog most closely related to the mammalian HP1 proteins [6, 11]. The goal of this study was to understand the basic function of HP1B, as HP1B, contrary to other HP1 proteins, is poorly characterized. We generated null-mutations in *HP1b* and show that *HP1b* is a non-essential gene. Despite its non-essential nature, HP1B is important for organism health, as loss leads to decreased offspring number in females and lowered activity levels in both larvae and adults. In addition, animals lacking HP1B have altered food intake and body composition. Combined with the phenotypic characterization, gene expression analysis of the *HP1b* mutants and a targeted metabolite analysis suggest that loss of HP1B results in altered metabolism in these animals. Thus, our study provides a novel link between an HP1 family protein and metabolism.

## MATERIALS AND METHODS

### Fly culture

All fly lines were cultivated on standard cornmeal-sucrose media [29] or Jazzmix Drosophila media (Fisher Scientific) at 25**°**C and 70% humidity. Animals other than those used for the fertility and PEV assays, were raised under controlled 12h light/ 12h dark cycle. The starting stock to generate the new *HP1b* alleles carrying the P{PTT-GB}HP1b^CB02849^ insertion was obtained from the Carnegie Protein Trap Library [30]. The insertion is located in the 5’ UTR of *HP1b*.

### Generation of *HP1b* mutant alleles

The *P* element of *HP1b^CB02849^* was mobilized by crossing homozygous females (*HP1b^CB02849-mw^, y^-^/HP1b^CB02849-mw^, y^-^*) to males containing the Δ2-3 *P* element transposase (*w*; ry^506^ Sb^1^ P{*Δ*2-3}99B/TM6B, Tb^1^*; Bloomington Drosophila Stock Center #1798). From the F1 population, males with the *HP1b^CB02849^* allele and the Δ2-3 *P* element transposase were selected (by the *Sb^1^* phenotype) and crossed to balancer females (*FM7j, B^1^, y^-^, w^-^/Df(1)ED6957^HP1b^, y^+^, w^mw^*; derived from Bloomington Drosophila Stock Center #8033). The *HP1b^CB02849^* transgene contains a copy of *mini-white* (*mw*), resulting in red eye pigmentation (*mw*).

Thus, putative *HP1b* mutants with the FM7j balancer (*HP1b*, y^-^/FM7j, B^1^, y^-^, w^-^)* among the progeny from the cross were identified based on the lack of red eye color and the presence of a *y^-^* phenotype, indicating the removal of the P{PTT-GB}HP1b^CB02849^ insertion. Putative mutants also were required to lack the Δ2-3 *P* element transposase by selecting against the *Sb^1^* phenotype. From 130 single male crosses set up in step 2, 89 female offspring lacking red eye color and with a *y^-^* phenotype were identified. Stocks were established, and males (HP1b*/Y) were screened by PCR (5’ – CTA GCG CAC GTG CAA CGC AAC – 3’; 5’ – ATA AAG GCG CTA GGT GGG CAG C −3’) to identify deletions. In addition to many small insertions, we identified three deletion alleles: *HP1b^16^* (Δ539bp), *HP1b^86^* (Δ1078bp), and *HP1b^49^* (Δ1309bp). Prior to carrying out the analyses described below, the *HP1b^16^* (Δ539bp) and *HP1b^86^* (Δ1078bp) alleles that are characterized in detail here were backcrossed into to a *y^-^w^-^* (*yw*) control strain for 6 generations to minimize genetic background effects.

### RT-PCR analysis

RNA was isolated from 2-4h embryos using Trizol according to the manufacturer’s recommendations (Invitrogen). The RNA was DNase treated and cDNA was prepared using Superscript III reverse transcriptase according to the manufacturer’s recommendations (Invitrogen). The cDNA was assayed for the presence of specific transcripts using PCR and the following primer combinations: *APC4* (nested PCR; primer set 1: 5’ – GTC ACC TTT CCC ACG CCC GG – 3’ and 5’ – CGT ACG GCT GGG AAT TGG CGT −3’; primer set 2: 5’ – GAC GCC GAG AGC GGA ACG AT – 3’ and 5’ – TGA GAT CGA TGC TGC CCG CC – 3’), *HP1b* (5’ – TCC GCG CAG CGA AAA CAC CT – 3’; 5’ – TAC CAT TGC CGC TGC CCG TG – 3’), *CG7426* (5’ – TGC AGC TTG CCC GTG AGC TC – 3’; 5’ – AGC GCC TGC TGC CGG AAT AC −3’), and *Rpl32* (positive control; 5’ – CGA TCT CGC CGC AGT AAA C – 3’; 5’ – CTT CAT CCG CCA CCA GTC G – 3’).

### Western blot

Nuclei were isolated from 20 flies of each genotype. Briefly, the animals were homogenized with a dounce homgenizer in lysis buffer (20mM HEPES, pH7.5; 10mM KCl; 3mM MgCl_2_; 0.2mM EGTA; 0.25mM sucrose; 0.5% NP-40; 1mM DTT; 1x protease inhibitor cocktail). The homogenate was filtered (Miracloth), and the nuclei pelleted by centrifugation (4°C; 15min.; 6000xg). The nuclei were washed twice with wash buffer (20mM HEPES, pH7.5; 10mM KCl; 3mM MgCl_2_; 0.2mM EGTA; 1mM DTT; 1x protease inhibitor cocktail), and then resuspended in RIPA buffer (50mM Tris-HCl, pH8.0; 2mM EDTA; 150mM NaCl; 0.5% Na deoxycholate; 0.1% SDS; 1% NP-40; 1mM PMSF; 1x protease inhibitor cocktail). Proteins were separated on 12% SDS-PAGE gels according to standard protocols and transferred to PVDF membrane. The Western blot was processed according to standard protocols. HP1B was detected using Q4111B (SDI; 1:2500); actin was detected using JLA20 (Iowa Hybridoma Bank; 1:100).

### Fertility assay - Female

Single virgin females (0-5 days old) from the *yw*, *HP1b^16^* and *HP1b^86^* strains were mated to two *yw* males in individual vials. After three days, these flies were moved to new vials, and again to a third set of vials on day six after the initial mating was set up. At day nine, the flies were discarded. From each of the three vials thus obtained from each female, offspring were counted four times, on days 10, 12, 14, and 16 after the vial was initially set. The offspring number per female was calculated by combining all offspring from the individual counts (12 total). Differences in offspring number between genotypes were analyzed by analysis of variance (ANOVA) and t-tests using R [31].

### Fertility assay - Male

Single virgin males (0-5 days old) from the *yw*, *HP1b^16^* and *HP1b^86^* strains were mated to two *yw* females in individual vials. After three days, these flies were moved to new vials, and again to a third set of vials on day six after the initial mating was set up. At day nine, the flies were discarded. From each of the three vials thus obtained from each male, offspring were counted four times, on days 10, 12, 14, and 16 after the vial was initially set. The offspring number per male was calculated by combining all offspring from the individual counts (12 total). Differences in offspring number between genotypes were analyzed by ANOVA and t-tests using R [31].

### Position effect variegation assay – *white* variegation

Eye pigment levels were measured by crossing males containing the *P* element reporter stock males to *yw, HP1b^16^, or HP1b^86^* homozygous virgin females. Offspring from the cross were aged to 3-5 days, and acid-extracted eye pigment was measured at OD_480_ for male and female offspring separately [adapted from [32]]. For each assay, five flies were used, with 6-10 biological replicates per genotype. Due to the location of *HP1b* on the X chromosome, male offspring from this cross lack wildtype *HP1b*, while females have one dose. Differences in eye pigment levels between genotypes were analyzed by ANOVA and t-tests using R [31].

### Position effect variegation assay – *Stubble* variegation

To assay the impact of *HP1b* mutations on Stubble variegation, the *Sb^V^* allele was used, similar to the assay described in Csink *et al* 1994 [33]. Males carrying the *Sb^V^* allele [*T(2,3)Sb^V^, ln(3)Mb, Sb^1^, sr^1^/TM3, Ser^1^*; Bloomington Stock # 878] were crossed to *yw, HP1b^16^, or HP1b^86^* homozygous virgin females. Offspring lacking the serrated wing phenotype associated with *Ser^1^* were examined, and the number of wildtype bristles was scored. In total, ten bristles per animal where assayed (four scutellar bristles, four dorsocentral bristles, and two posterior post-alar bristles), and the number of wildtype bristles was recorded for each animal. Due to the location of *HP1b* on the X chromosome, male offspring from this cross have lack wildtype *HP1b*, while females have one dose. Differences in bristle number between genotypes were analyzed by ANOVA, Tukey’s HSD *post hoc* tests, and t-tests using R [31].

### Animal activity – third instar larvae

Sets of ten third instar larvae were placed in the center of a petri dish containing 1.5% agar. The petri dish was placed onto a diagram of concentric circles spaced 1cm apart to evaluate the movement of the larvae. The larvae were allowed to move freely, and pictures were taken with a digital camera every 15 seconds for 2 minutes. From the images, the number of animals in each concentric circle were recorded to provide an estimate of the distance traveled by the animals. For each genotype (*yw, HP1b^16^, or HP1b^86^*), five trials with eight larvae each were performed. The data from the 60 second time point were recoded to divide all animals into two categories: those animals that remained in the center of the petri dish (“no movement”) and those animals that left the center of the petri dish (“movement”). The fraction of animals moving versus not moving was compared between the *HP1b* mutant and *yw* control genotypes (chi-square test) using R [31].

### Animal activity – adult flies

Basal activity of the animals was monitored using a locomotion activity monitor (LAM25) from TriKinetics. 3-5 day old virgin flies from the *yw, HP1b^16^*, and *HP1b^86^* strains were placed into assay vials (n= 5 for each sex and genotype; ten flies per vial). Next, the vials were placed into the monitor, and the monitor was placed in the incubator (25°C, 60% humidity, 12h light-dark cycle). The flies were allowed to recover for 1h prior to the start of the assay period. Animal activity was recorded every 5 minutes from 8AM to 10AM. We performed five independent replicates of this assay. Total activity during the 2hr period was used in the analyses. ANOVA (factors: sex, genotype, trial) and Tukey’s HSD *post hoc* tests were performed in R to investigate the sources of the variation observed between the samples [31].

### Stress assay – starvation

3-5 day old virgin flies were moved onto starvation media (1.5ml of 1% agar to provide hydration) and placed into an incubator (25°C, 60% humidity, 12 hr light-dark cycle). Deaths were recorded every 8 hours until all flies had died (n= 100 for each sex and genotype [*yw*, *HP1b^16^*, *HP1b^86^*]; 20 flies per vial) [34]. Three independent replicates of this assay were performed. Survival analysis was carried out in R [31], using the “Survival” package, specifically its survival difference formula (Kaplan-Meier), to compare the survival curves between the three genotypes [31, 35].

### Stress assay - Heat stress

3-5 day old virgin flies were transferred to fresh Jazzmix medium (Fisher Scientific) and placed in a warm room maintained at 37°C. Flies were monitored every hour until all flies had died (n= 100 for each sex and genotype; 20 flies per vial) [36]. We performed three independent replicates of this assay. Survival analysis was carried out in R [31], using the “Survival” package, specifically its survival difference formula (Kaplan-Meier) [31, 35].

### Stress assay – Oxidative stress

To assay oxidative stress, flies were exposed to the herbicide paraquat, an oxidizer. Filter paper disks saturated with paraquat solution (20mM paraquat, 5% sucrose) were placed into vials containing starvation media (1.5ml of 1% agar). 3-5 day old virgin flies were placed inside vials (n= 100 for each sex and genotype; 20 flies per vial), returned to the incubator (25°C, 60% humidity, 12 hr light-dark cycle), and monitored every 4 hours until all flies had died [37]. We performed three independent replicates of this assay. Survival analysis was carried out in R [31], using the “Survival” package, specifically its survival difference formula (Kaplan-Meier) [31, 35].

A second oxidative stress assay was carried out to determine if the increased survival of *HP1b* mutants was related to their lower food and thus paraquat intake. In this assay, the paraquat concentration was scaled based on food consumption (Figure 7A); thus, the paraquat concentration for *yw* was decreased to 13.2mM paraquat. We performed this assay with 100 animals for each sex and genotype, again with 20 flies per vial. Data analysis was performed in R [31] using the R package “Survival,” specifically, its survival difference formula (Kaplan-Meier) [31, 35].

### Lifespan Assay

To determine lifespan, virgin flies were maintained on Jazzmix (Fisher Scientific) under standard conditions (25°C, 60% humidity, 12 h light-dark cycle). Flies were moved to new food every three days without the use of CO_2_ and monitored daily until all flies had died (starting population: n= 100 for each sex and genotype; 20 flies per vial) [36, 38]. We performed three independent replicates of this assay. Survival analysis was carried out in R [31] using the “Survival” package, specifically its survival difference formula (Kaplan-Meier) [31, 35].

### Food Consumption assay

Capillary Feeder (CAFÉ) assays were carried out as previously described [39]. Briefly, for the assay, animals were placed on starvation media (1.5ml of 1% agar to provide water) in standard fly vials that were cut in half, with four equally spaced holes around the perimeter. A nutrient solution (5% sucrose, 5% yeast solution) was provided through two capillaries inserted into the foam vial top. At 4PM of the day prior to the start of the assay, 3-5 day old virgin flies were placed into the assay vials (five flies per vial; n= 4-8 for each sex and genotype), the capillaries were filled with the nutrient solution, and the flies were returned to the incubator (25°C, 60% humidity, 12 hr light-dark cycle) to acclimatize overnight to the new feeding method. At 8AM, the capillaries were refilled with the nutrient solution. Eight hours later, the drop in liquid levels within the capillaries was measured. We assayed the same flies twice over two eight-hour time periods (8AM to 4PM) on two consecutive days, and used average food consumption for further analysis. Evaporation was estimated using control vials lacking flies. We performed three independent replicates of this assay. Due to differences in the capillaries used, the three trials were analyzed separately. Data analysis was performed in R using ANOVA with a Tukey’s honest significant difference (HSD) *post hoc* test and Kruskal-Wallis tests in case of non-normality [31].

### Body Composition Analysis

Body fat content was measured using quantitative magnetic resonance (QMR) for flies at specific ages at the UAB Small Animal Phenotyping Core. Virgin flies were collected, sorted by sex and genotype, and aged for the desired time (3-5, 20, 30, 40 or 50 days old). At the appropriate age, flies were placed into a microcentrifuge tube (10 flies per tube, 4 tubes per sex and genotype for each time point), and stored on ice. QMR analysis was performed by the UAB Small Animal Phenotyping core with an EchoMRI 3-in-1 machine (Echo Medical Systems, Houston, TX). Flies were transferred into the biopsy tube and scanned using the biopsy setting with 9 primary accumulations. Data were analyzed in R using ANOVA with a Tukey’s HSD *post hoc* test [31].

### RNA-seq analysis

RNA was isolated using Trizol from third instar larvae of the *HP1b^16^* and *HP1b^86^* mutant stocks as well as of the *yw* control strain. Integrity of the RNA samples was confirmed by formaldehyde agarose gel electrophoresis. The RNA samples were prepared for RNA-seq analysis by the UAB Heflin Center for Genomic Science Genomics Core Lab. 30-40 million RNA-seq reads were collected per sample using the Illumina Sequencing Platform, and two samples per genotype were processed. The UAB Heflin Center for Genomic Science Genomics core processed the raw data using the Tuxedo suite software (Bowtie, Tophat, Cufflinks) and identified genes differentially expressed between the *HP1b* mutants and the *yw* control strain by Cuffdiff [40]. Only genes meeting an FDR (false discovery rate) cut-off of 0.05 were considered in the downstream analyses.

### GO term enrichment analysis

The RNA-seq analysis revealed that while neither *HP1b* allele generated full-length mRNA or functional HP1B protein, an aberrant RNA was produced from the *HP1b* locus in the *HP1b^16^* stock. To minimize the potential impact of this aberrant mRNA on the interpretation of results, only genes misregulated in both *HP1b* mutant strains were included in the GO term analysis. Differentially expressed genes were classified as up-versus down-regulated, and genes misregulated in both *HP1b* mutant strains were identified using custom scripts. GO term enrichment analysis was carried out using PANTHER [41].

### Krebs cycle metabolite analysis

Metabolite concentrations were measured by mass spectrometry for virgin flies aged 3-5 days. Flies were sorted by sex and genotype, aged 3-5 days, flash frozen in liquid nitrogen, homogenized in cold methanol, and delivered to the UAB Targeted Metabolomics and Proteomics Laboratory (TMPL) for analysis (n=4 biological replicates per sex and genotype). At the TMPL, supernatants were stored at −80°C until further processing. Standards were generated as a master mix of all compounds at 100 μg/mL in water and serial diluted to 10x of the final concentrations (0.05-10 μg/ml, 9 standards). Standards were further diluted to 1x in methanol to a total volume of 1 mL, and dried by a gentle stream of N_2_. For cell extracts, 1 mL of each were transferred to a glass tube and dried under a gentle stream of N_2_. Standards and samples were resuspended in 50μl of 5% acetic acid and vortexed for 15 seconds. Amplifex™ Keto Reagent (SCIEX, Concord, Ontario, Canada) (50 μL) was added to each sample and allowed to react for 1h at room temperature. Standards and samples were then dried under a gentle stream of N^2^ and resuspended in 1ml of 0.1% formic acid. Samples were analyzed by liquid-chromatography(LC)-multiple reaction ion monitoring-mass spectrometry. Liquid chromatography was performed using a LC20AC HPLC system (Shimadzu, Columbia, MD) with a Synergi Hydro-RP 4 μm 80A 250 x 2 mm ID column (Phenomenex, Torrance, CA). Mobile phases were: A) 0.1% formic acid and B) methanol/0.1% formic acid. Compounds were eluted using a 5-40% linear gradient of B from 1 to 7 min, followed by a column wash of 40-100% B from 7 to 10 min, and re-equilibrated at 5% B from 10.5 - 15 min. Column eluant was passed into an electrospray ionization interface of an API 4000 triple-quadrupole mass spectrometer (SCIEX). The following mass transitions were monitored in the positive ion mode: m/z 261/118 for α-ketoglutarate, m/z 247/144 for oxaloacetate and m/z 204/144, 204/118 and 204/60 for pyruvate. In the negative mode, the following transitions were monitored: m/z 115/71 for fumarate, m/z 89/43 for lactate, m/z 117/73 for succinate, m/z 133/115 for malate, m/z 173/85 for cis-aconitate, m/z 191/87 for citrate, m/z 191/73 for isocitrate, m/z 147/129 for 2-hydroxyglutarate, m/z 146/102 for glutamate, m/z 145/42 for glutamine and m/z 132/88 for aspartate. The 16 transitions were each monitored for 35ms, with a total cycle time of 560ms. MS parameters were CAD 4, CUR 15, GS1 60, GS2 30, TEM 600, IS −3500 volts for negative polarity mode and IS 4500 for positive polarity mode. Peak areas of metabolites in the sample extracts were compared in MultiQuant software (SCIEX) to the those of the known standards to calculate metabolite concentrations. MetaboAnalyst 3.0 was used for the analysis [42], with additional data analysis performed in R [31].

### Mitochondrial complex analysis

Mitochondrial complex 3 and citrate synthase activity were measured utilizing the resources at the UAB Bio-analytical Redox Biology (BARB) core. Thoraces were dissected from 3-5 day old male animals (five biological replicates per genotype) and placed in isolation buffer on ice [43]. The thoraces were homogenized, remaining solids removed by filtration, and protein concentration was measured with a Lowry assay [44]. To assay citrate synthase, a coupled reaction with acetyl-CoA, oxaloacetate, and 5,5-dithiobis-2-4 nitrobenzoic acid was used, and the results were measured using a spectrophotometer [45, 46]. Complex III activity was measured by assaying the conversion of cytochrome C (its electron acceptor) from the oxidized to the reduced form using a spectrophotometer to detect a shift in absorbance [47]. Data analysis was performed in R [31].

### Data and reagent availability

Drosophila strains are available upon request for a minimum of three years after the publication date. Raw data files for all analyses are available as a supplemental file (compressed archive). RNA-seq data sets are available from GEO (accession number xxx).

## RESULTS

### *HP1b* is a non-essential gene

We generated loss-of-function mutations in the HP1 family gene *HP1b* by imprecise *P* element excision to study its function. Our screen generated three deletion alleles, *HP1b^16^*, *HP1b^86^*, and *HP1b^49^* (Figure 1A). The deletions span 539bp, 1,078bp, and 1,309bp respectively, with *HP1b^16^* and *HP1b^86^* deleting major portions of the *HP1b* coding sequence, and the *HP1b^49^* deletion extending into the intergenic region between *HP1b* and its downstream neighbor *APC4* (*Anaphase Promoting Complex 4*). *HP1b^16^* and *HP1b^86^* are homozygous viable, while *HP1b^49^* is homozygous lethal early in development (no homozygous first instar larvae are observed). As *HP1b^16^* and *HP1b^86^* both lack full-length *HP1b* mRNA and protein (Figure 1B and 1C), we attribute the lethality of *HP1b^49^* to an effect on *APC4* due to the deletion of some of the regulatory sequences presumably present in the intergenic sequences between *HP1b* and *APC4*. While the full-length *HP1b* mRNA is absent in *HP1b^16^* and *HP1b^86^* mutant embryos, the two neighboring genes *APC4* and *CG7426* are expressed (Figure 1B). Thus, our analysis here focuses on the *HP1b^16^* and *HP1b^86^* alleles. At 25**°**C, homozygous stocks of *HP1b^16^* and *HP1b^86^* are viable and can be maintained without difficulties under standard conditions. Thus, we conclude that *HP1b* is a non-essential gene in contrast to the centromeric HP1 family member, HP1a, encoded by *Su(var)205*, which is homozygous lethal at the third instar stage [24, 48].

**Figure 1.**
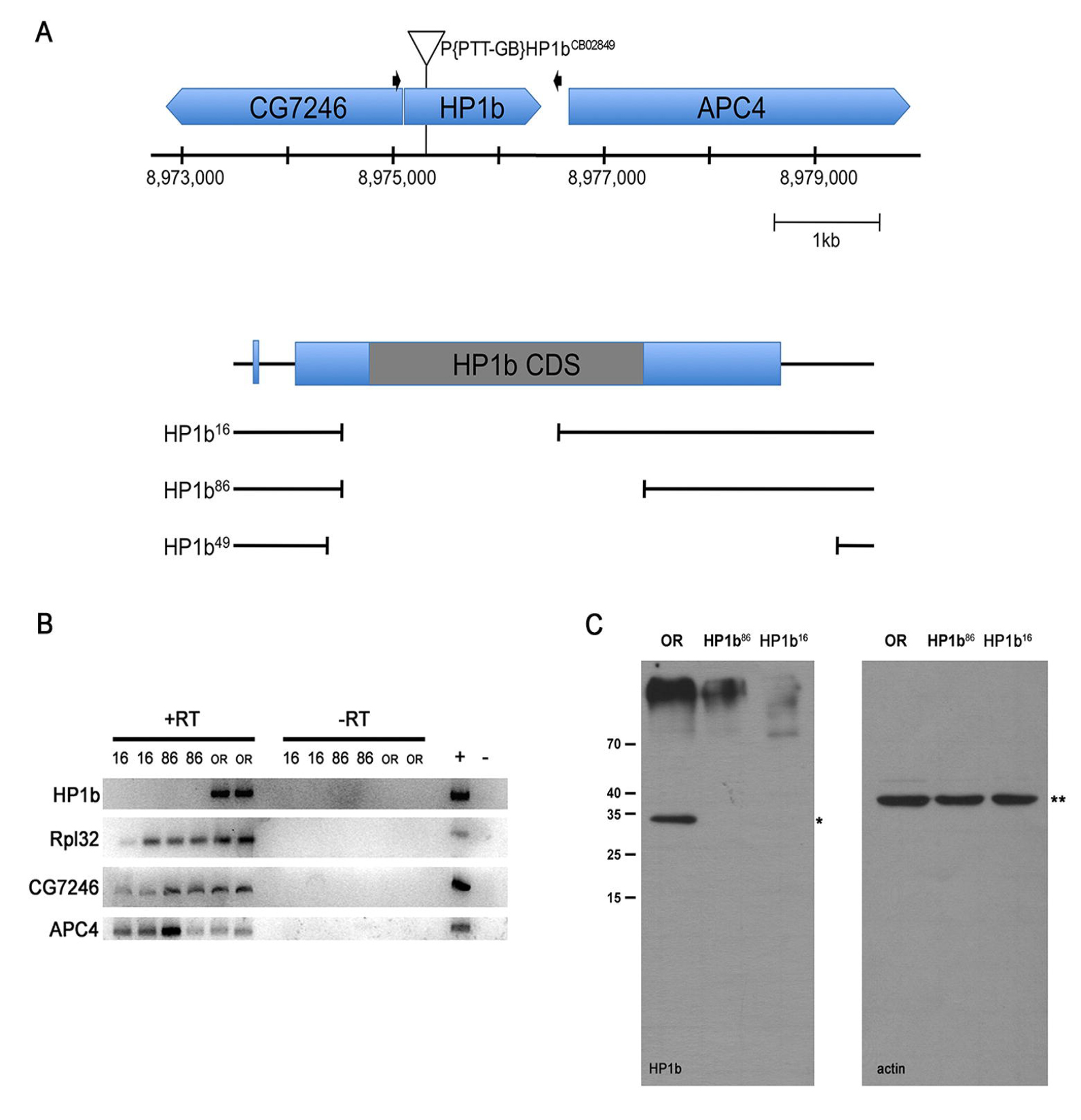
*HP1b^16^* and *HP1b ^86^* are null alleles. **A.** Top - diagram showing the genomic region near the *HP1b* locus on the X chromosome. The *P* element mobilized to generate the *HP1b* alleles is indicated by a triangle above the genes shown in blue above the position on the X chromosome (bp). Black arrows mark the primers used to screen for deletions. Bottom – close-up of *HP1b* with the three deletion alleles shown below. All three deletions remove the *HP1b* start codon. **B.** RT-PCR analysis shows that full-length *HP1b* mRNA is transcribed in wildtype OR embryos, but not in *HP1b^16^* or *HP1b^86^* homozygous embryos. *CG7346* and *APC4* are transcribed in all samples, as is the *Rpl32* positive control. RT: reverse transcriptase. **C.** No HP1B protein is detected in *HP1b^16^* and *HP1b^86^* homozygous larvae (left panel; *); Actin (right panel; **) serves as loading control.

### Mutations in *HP1b* impact the number of offspring produced by females

Expression data generated by modENCODE available from Flybase (www.flybase.org, [49]) indicate that expression of *HP1b* is moderate throughout development with high expression during early embryonic development (2-4hr) and in the ovaries [50, 51], suggesting that *HP1b* might impact the number of offspring produced per female. To investigate this hypothesis, we assayed the potential impact of *HP1b* mutations on offspring number. When the average offspring number of single females mated to two males of *y^-^w^-^* (*yw*) is compared between *HP1b* mutants and control *yw* females, a reduction in offspring number is observed for the mutants (Figure 2A). While the *yw* females had on average 56 offspring, *HP1b^16^* and *HP1b^86^* had 31 and 36 offspring on average respectively, a drop that is statistically significant (p<0.01 for *HP1b^16^* and p<0.05 for *HP1b^86^*; t-test). As a control, we also measured the average number of offspring produced by individual *HP1b* and *yw* males mated to two *yw* females. We found no significant differences in male offspring number between the *HP1b* mutant animals and the *yw* control strains (Figure 2B, p-value not significant). These data indicate that the lack of HP1B protein specifically impacts the offspring number of female animals.

**Figure 2.**
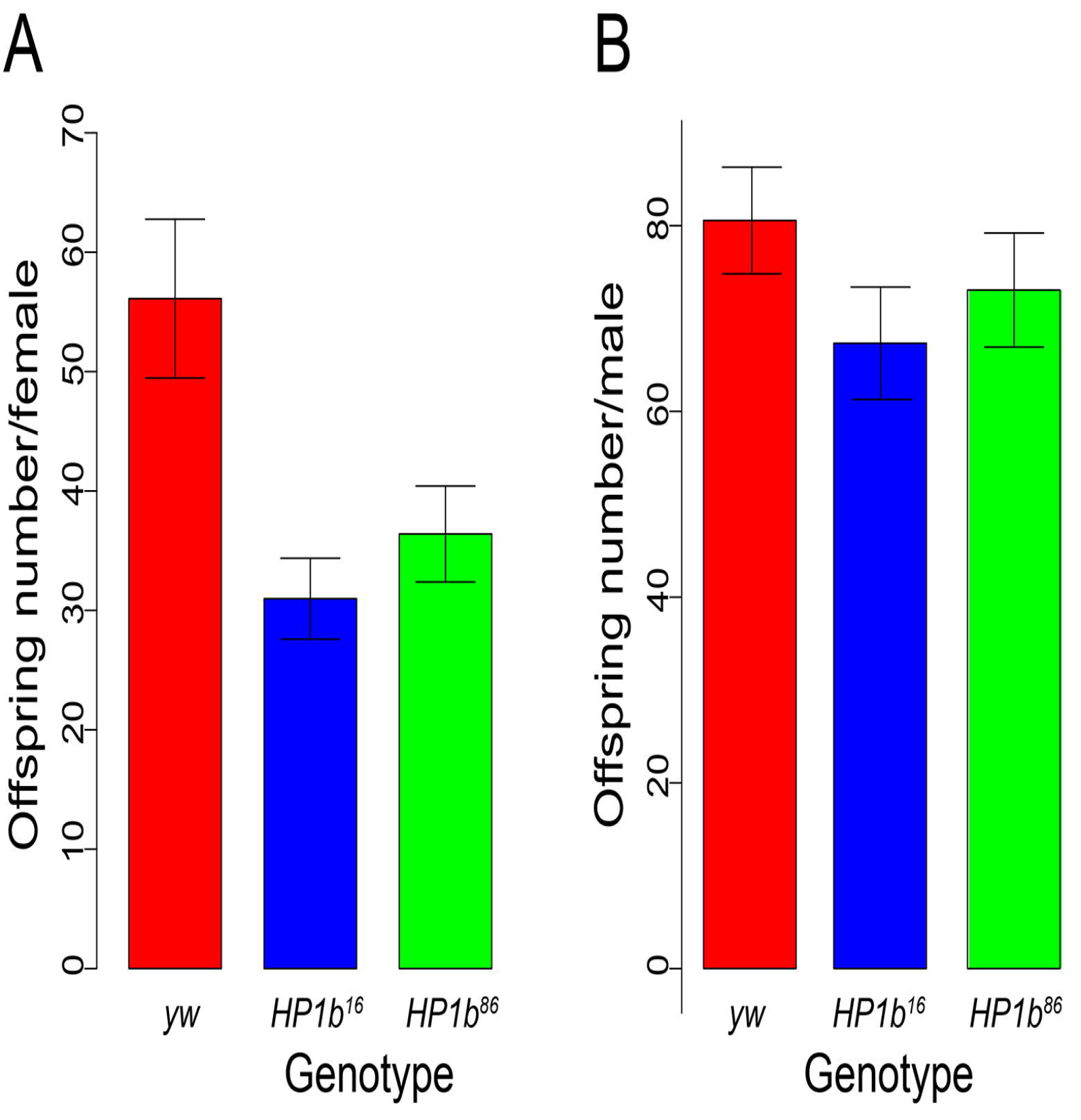
Mutations in *HP1b* reduce female fertility. **A.** The average offspring number for individual females of *yw* (red; n=73), *HP1b^16^* (blue; n=80), and *HP1b^86^* (green; n=82) is shown (Y-axis). Error bars: SEM. The average number of offspring in both *HP1b^16^* and *HP1b^86^* females is significantly reduced compared to *yw* females (t test, p<0.05). **B.** The average offspring number for individual males of *yw* (red; n=80), *HP1b^16^* (blue; n=73), and *HP1b^86^* (green; n=76) is shown (Y-axis). Error bars: SEM. The average offspring number of *HP1b^16^* and *HP1b^86^* males does not differ from that of *yw* males (t test, not significant).

### *HP1b* is an Enhancer of Variegation

Position effect variegation (PEV) analysis in Drosophila is a powerful tool to investigate the impact of mutations on transgene expression in particular chromatin environments. Transgene expression from reporters within or close to heterochromatin tends to variegate due to silencing of the reporter in some cells but not others. If a mutation for example affects the chromatin structure, increased expression or suppression of variegation [Su(var) phenotype] is seen if the protein is required for heterochromatin formation or has repressive functions. If on the other hand decreased expression or enhancement of variegation is seen [E(var) phenotype], this result indicates that the protein is involved in counteracting heterochromatin formation or has activating functions. This assay has been used successfully to demonstrate that mutations in HP1a increase expression of a variety of reporter genes [Su(var) phenotype] [48], indicating it can act as a transcriptional repressor. In contrast, tethering HP1C to a reporter gene increased its expression, indicating that it functions as a transcriptional activator [52]. We used PEV analysis to investigate the role of HP1B in the regulation of gene expression. Males containing a *P* element reporter that fuses the *hsp70* promoter to the *white* coding region were crossed to virgin females from the homozygous *HP1b^16^* and *HP1b^86^* stocks as well as to *yw* as a control. As PEV effects often depend on the specific reporter insertion used, five different reporter lines were tested: 39C-2 (chr. 2; pericentric), 39C-3 (chr. 2, pericentric), 118E-10 (chr. 4; pericentric), 39C-12 (chr. 4; arm), and 39C-27 (telomeric) [53]. Assaying males, which lack HP1B entirely due to the gene’s location on the X chromosome, we find that eye pigment levels are reduced in the presence of the *HP1b* mutations compared to the *yw* control (Figure 3A). This E(var) phenotype is consistent for all five reporters assayed, including pericentric and telomeric reporter insertions (Figure 3B). In females, which are heterozygous for the *HP1b* mutations, the results differ depending on the reporter location and include E(var) responses (39C-2 and 39C-3) as well as Su(var) responses (118E-10 and 39C-12) (Figure S1A). Thus, *HP1b* mutations act as modifiers of PEV in both sexes for these *hsp70-white* reporters, and they have a strong E(var) effect in males in all genomic contexts assayed.

**Figure 3.**
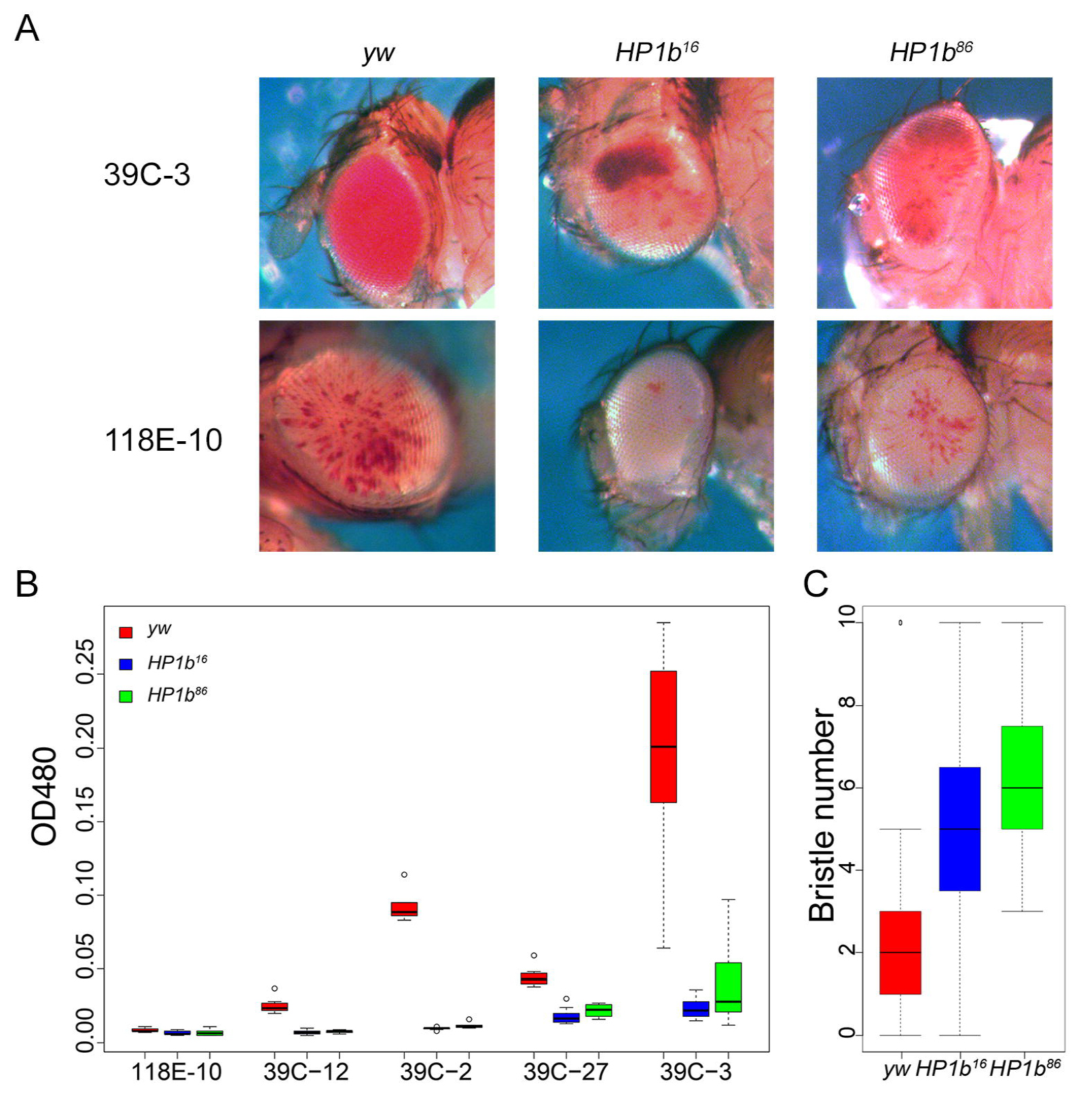
*HP1b* is an enhancer of variegation. **A.** If mutant *HP1b* alleles are introduced into stocks carrying *hsp70-white* reporter genes, the red eye color (reflecting *white* expression) is decreased compared to the *yw* control in male flies. Representative eye pictures are shown from males for two of the reporter insertions, 39C-3 located in the pericentric region of chromosome 2, and 118E-10 located in the pericentric region of chromosome 4. **B.** Quantitative eye pigment assays for males from the five reporter insertions assayed. All comparisons of mutant eye pigment values to the *yw* control are statistically significant (p<0.05, t-test). X-axis: reporter insertions. Y-axis: OD_480_ measuring eye pigment. Results from the *yw* control are in red, from *HP1b^16^* in blue, and from *HP1b^86^* in green. Box plots: black bar - median; box - +25% and −25% quartiles; whiskers – maximum and minimum; circles – outliers; n=6-10. **C.** *Stubble* variegation assays indicate that *HP1b* is an enhancer of variegation, as the presence of the *HP1b* mutant alleles significantly increases the number of wildtype bristles compared to the *yw* control (p<0.001, Tukey multiple comparisons of means; n=19-39). Data from males. Results from the *yw* control are in red, from *HP1b^16^* in blue, and from *HP1b^86^* in green. Box plots: black bar - median; box - +25% and −25% quartiles; whiskers – maximum and minimum; circles – outliers. For data from females, see Figure S1.

To ensure that the E(var) effect upon complete loss of HP1B is a general phenomenon and not restricted to the *white* reporter used in these assays, we investigated a second PEV system, variegation of the *Stubble* (*Sb)* allele *Sb^V^*. *Sb^V^* is a gain-of-function allele that causes a bristle defect. Animals heterozygous for *Sb^V^* have a mixture of wildtype and defective bristles. If an E(var) mutation is crossed to the *Sb^V^* reporter, the fraction of defective bristles will decrease, while in the presence of a Su(var) mutation, the fraction of defective bristles will increase. Males containing the *Sb^V^* reporter were crossed to virgin females from the homozygous *HP1b^16^* and *HP1b^86^* stocks as well as to *yw* as a control. In the offspring, ten bristles (four scutellar bristles, four dorsocentral bristles, and two posterior post-alar bristles) were examined, and the number of wildtype bristles was recorded. We find that the presence of *HP1b* null alleles significantly increases the number of wildtype bristles compared to the *yw* control (Figure 3C; Tukey’s test, p<0.01). In males, for example, the wildtype bristle number increased approximately 2-fold from an average of 2.7 wildtype bristles per animal in *yw* to 5.3 and 6.1 in the *HP1b^16^* and *HP1b^86^* animals respectively. *HP1b* mutations also increase the wildtype bristle number in females (Figure S1B), and thus, *HP1b* mutations generally act as enhancers of variegation for the *Sb^V^* reporter. These findings confirm that the results obtained with the *hsp70-white* reporters and suggest that complete loss of HP1B results generally in enhancement of variegation as indicated by the effects in diverse genomic contexts with multiple reporters.

### *HP1b* mutants have decreased activity levels

During the experiments investigating PEV and offspring number, we noticed that the *HP1b* mutants tended to be easier to handle than the *yw* controls as they appeared to move less. To determine if mutations in *HP1b* indeed affected the activity levels of the animals, experiments were carried out in third instar larvae and adult flies. In third instar larvae, we assayed crawling ability and determined the fraction of animals that moved or failed to move in a 60 second time period (Figure 4A). In this assay, *HP1b* mutant larvae tended to remain unmoving more often. For example, only 21% of the larvae from the *HP1b^16^* strain moved during the assay period (10/47), while 48% of the *yw* larvae moved (23/48). Altogether, significantly fewer *HP1b* mutant animals moved away from the center of the assay dish during the assay period (p< 0.001 for both mutant alleles, chi-square test, Figure 4A). These data indicate that lack of HP1B leads to lower activity levels in third instar larvae.

**Figure 4.**
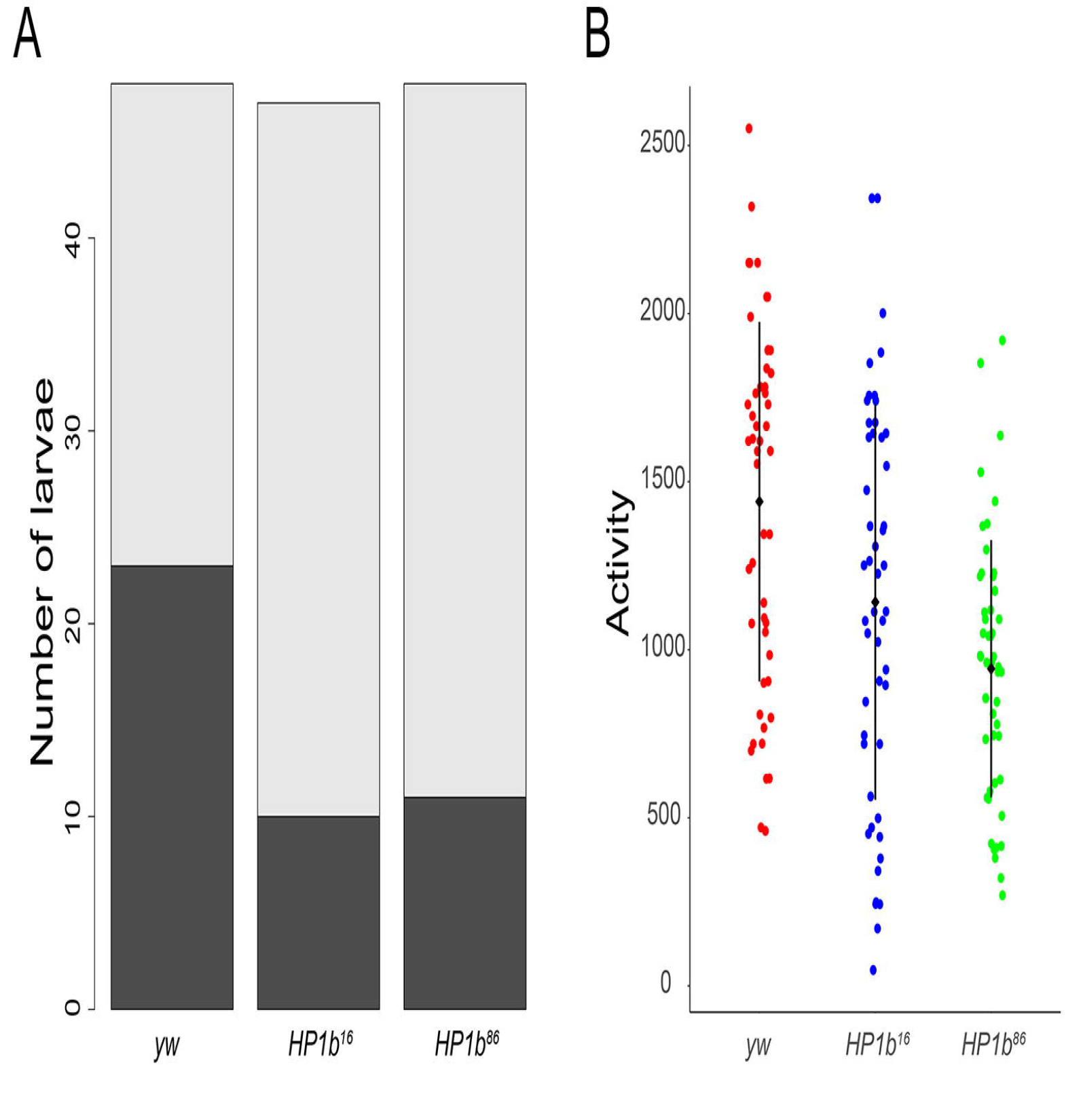
Animals lacking HP1B show decreased levels of activity. **A.** A significantly higher fraction of third instar larvae lacking HP1B (*HP1b^16^* and *HP1b^86^*) failed to move in a 60 second observation period compared to the *yw* control (p<0.001 for both comparisons; chi-square test). Dark grey: animals that moved; light grey: animals that failed to move. Y-axis: Number of larvae. N=47-50 per genotype. **B.** Adult animals lacking HP1B show significantly lower activity levels than animals of the *yw* control genotype (p<0.001 for both comparisons; Tukey’s HSD). Y-axis: activity level as measured by recorded beam crossings in an activity monitor for 10 flies within a 2hr assay period. Black diamond and bar: mean +/- SD. N=50 per genotype.

Next, activity levels of adult animals were assayed to determine if the lower activity level observed in larvae is a general feature of *HP1b* mutants consistent across different developmental stages. We utilized an activity monitor which measures fly activity levels by recording every instance a fly crosses the middle of the assay chamber. In this assay, *HP1b* mutants have significantly decreased levels of activity compared to the controls (Figure 4B). In a two-hour period, an average of 1142 crossings were recorded for *HP1b^16^* mutants, *HP1b^86^* mutants crossed the middle of the assay chamber on average 944 times, while the *yw* control flies logged 1441 crossings (Figure 4B). These differences are highly significant (Tukey’s HSD; p<0.001 for both mutant alleles). While activity levels are significantly different between males and females (ANOVA, p<0.001), the decrease in activity associated with mutations in *HP1b* is observed in both males and females (Figure S2). However, the decrease is more pronounced in females, where the decrease is significant for both *HP1b^16^* and *HP1b^86^* (Tukey’s HSD, p<0.001), while in males, the activity of only *HP1b^86^* mutants is significantly decreased compared to *yw* (Tukey’s HSD, p<0.001). Taken together with the results from the larval crawling assay, these results suggest that lowered animal activity levels are a general feature of *HP1b* mutants, irrespective of developmental stage.

### Loss of HP1B alters stress resistance

Given that loss of HP1B impacts the number of offspring produced by the animals as well as their activity levels, we questioned whether other aspects of the animals’ physiology might be impacted as well. First, we investigated stress resistance. In Drosophila, increased stress resistance is often observed in animals with increased lifespan [37, 54-57], and as stress assays require much less time, they are typically carried out prior to investigating lifespan. Three separate stress resistance assays were used to investigate resistance to starvation stress, oxidative stress, and heat stress. To assay resistance to starvation stress, flies were grown under standard conditions and transferred to vials containing 1% agar, providing hydration but no nutritional value. Deaths were recorded every eight hours, and survival curves were generated (Figure 5A). Survival curves of the *HP1b* mutant stocks are shifted to the right compared to the *yw* control, indicating increased stress resistance and longer survival.

**Figure 5.**
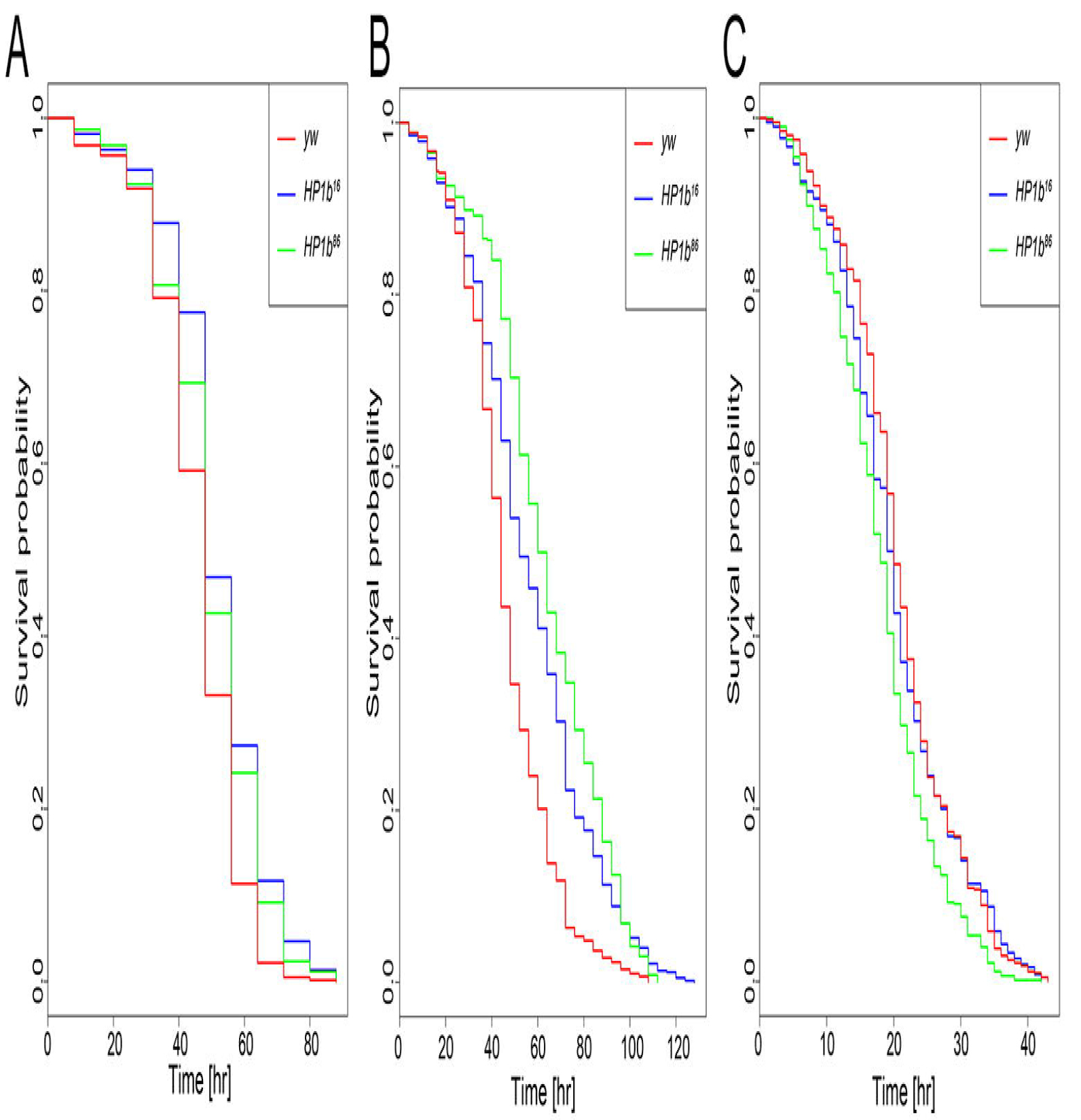
Loss of HP1B increases survival during starvation and oxidative stress, but not during heat stress. **A.** *HP1b* mutant animals survive significantly longer during starvation condition than *yw* control animals (p=1.022e-13 and p=7.188e-06 for *HP1b^16^* and *HP1b^86^*, respectively, Kruskal-Wallis rank sum test). **B.** *HP1b* mutant animals survive significantly longer after exposure to the oxidizer paraquat than *yw* control animals (p = 3.057e-13 and p< 2.2e-16, for *HP1b^16^* and *HP1b^86^*, respectively, Kruskal-Wallis rank sum test). **C.** *HP1b* mutant animals do not show improved survival during heat stress conditions (37°C) compared to *yw* control animals. In contrast, they show lower survival (p = 0.04158 and p = 1.874e-09, for *HP1b^16^* and *HP1b^86^*, respectively, Kruskal-Wallis rank sum test). For **A-C**: X-axis: Time to death in hours. Y-axis: Survival probability. Data shown are from three trials, each with 100 animals per genotype/sex. For separate survival curves for males and females, see Figure S3.

Living on average 51.65 and 49.37 hours under starvation conditions, *HP1b* mutants show significantly increased survival compared to *yw*, which lives an average of 45.60 hours (p=1.022e-13 and p=7.188e-06 for *HP1b^16^* and *HP1b^86^*, respectively, Kruskal-Wallis rank sum test). In addition, the survival curves are significantly different between *HP1b* mutant and *yw* control flies (Kaplan-Meier survival function: p=1.11e-16 for *HP1b^16^*, and p=9.83e-09 for *HP1b^86^*). The increased survival under starvation conditions was observed also when the data from males and females were analyzed separately (Figure S3A). In females, both the survival curves and means are different between *HP1b* mutants and *yw* (Kruskal-Wallis rank sum test: p=6.455e-09 and p=4.672e-07 for *HP1b^16^* and *HP1b^86^* respectively; Kaplan-Meier survival function: p=2.36e-10 for *HP1b^16^*, and p=4.67e-08 for *HP1b^86^*), while in males, the comparison of means is only significant for *HP1b^16^* (Kruskal-Wallis rank sum test: p=2.054e-07), despite significantly shifted survival curves for both *HP1b* strains (Kaplan-Meier survival function: p=1.86e-09 for *HP1b^16^* and p=0.011 *HP1b^86^*). Together, these data demonstrate that loss of HP1B allows flies to survive longer under starvation conditions.

Next, we investigated resistance to oxidative stress using a paraquat feeding assay. Paraquat is an herbicide that induces oxidative stress when ingested, and it is widely used to measure oxidative stress resistance in flies [37]. Flies were fed a paraquat-laced sucrose solution and monitored to record time of death. Results from this assay revealed that the survival curve of the *HP1b* mutants were shifted to the right, and the control flies (red) showed a steeper drop-off in survival rate than the two *HP1b* mutant strains (blue and green, Figure 5B). In the combined analysis of males and females, *HP1b* mutants have significantly increased average lifespan compared to the *yw* control strain: *HP1b^16^* live on average 56.39 hours under oxidative stress, *HP1b^86^* live 62.84 hours, and *yw* animals live 46.12 hours (p=3.057e-13 for *HP1b^1^*^6^ and p<2.2e-16 for *HP1b^86^*; Kruskal-Wallis rank sum test). In addition, the survival curves of *HP1b* mutants differ significantly from those of the *yw* control (Figure 5B; p=0 for *HP1b^16^* and *HP1b^86^*; Kaplan-Meier survival function). The same shift towards increased survival of the *HP1b* mutant strains is also evident when data from females and males are analyzed separately (Figure S3B; females: p=8.13e-13 for *HP1b^16^* and p=0 for *HP1b^86^*; males: p=6.96e-9 for *HP1b^16^* and p=1.29e-14 for *HP1b^86^*), and the average survival of the *HP1b* mutants is increased compared to *yw* (Figure S3B; females: p=8.16e-13 for *HP1b^16^* and p=0 for *HP1b^86^*; males: p=6.96e-09 for *HP1b^16^* and p=1.29e-14 for *HP1b^86^*). Together, these data show that loss of HP1B increases oxidative stress resistance as measured by paraquat feeding assays.

Lastly, resistance to heat stress was investigated. Flies were raised under standard conditions and then moved to 37°C. The flies’ time of death was recorded hourly to generate survival curves. In contrast to the findings for starvation and oxidative stress, no increased resistance to heat stress was detected in the *HP1b* mutants, neither in males nor in females (Figure 5C and Figure S3C). In contrast, the *HP1b* strains actually showed decreased survival compared to the control, with *HP1b^16^* surviving on average 20.04 hours, *HP1b^86^* surviving 18.01 hours, and *yw* surviving 20.80 hours (p=0.04158 for *HP1b^1^*^6^ and p=1.874e-09 for *HP1b^86^*; Kruskal-Wallis rank sum test). When males and females are analyzed separately, only the *HP1b^86^* strain shows significantly decreased survival, while *HP1b^16^* is not different from *yw* (females: p=0.001803; males: p=1.499e-08; Kruskal-Wallis rank sum test). Analysis of the survival curves shows similar results: only *HP1b^86^* has a survival curve that is significantly different from *yw*, be that in the combined analysis or in males/females (combined: p=2.93e-09; females: p=0.0002: males: p=7.02e-7; Kaplan-Meier survival function). These results indicate that *HP1b* mutant flies do not have increased heat stress resistance. This result is unexpected, because most often, stress resistance in general is increased, irrespective of assay. Thus, *HP1b* mutants are unusual in showing resistance to starvation stress and oxidative stress but not showing resistance to heat stress.

### Loss of HP1B impacts survival

In Drosophila, stress resistance phenotypes are often seen in animals that show increased longevity. Given the unusual stress resistance phenotypes we observed in the *HP1b* mutant strains, next, we investigated their longevity by generating survival curves. In three replicate experiments, we followed cohorts of 100 adult flies for each genotype/sex from hatching to death (Figure 6). Comparing the survival curves from all three strains, a statistically significant impact of “genotype” as a factor was detected (p=0.0483, log rank test) when the data from both sexes is combined for the analysis. This difference is mainly driven by the results from females, which show a strong effect of genotype (p= 0.00273, log rank test), which is absent in males (Figure S4). In the *HP1b* mutant strains, especially female animals experience less mortality early in life, until approximately day 40, when mortality rates increase. Towards the last quarter of the lifespan (~day 60), mortality rates in the *yw* control lines are lower than in the *HP1b* mutant strains, leading to a longer maximum lifespan (p=1.11e-15 for *HP1b^16^* and p=0 for *HP1b^86^*, log rank test on the last 10% of surviving animals). These results illustrate that loss of *HP1b* has a complex effect on survival, affecting the two sexes differently, and mainly impacting survival in early/mid-life.

**Figure 6.**
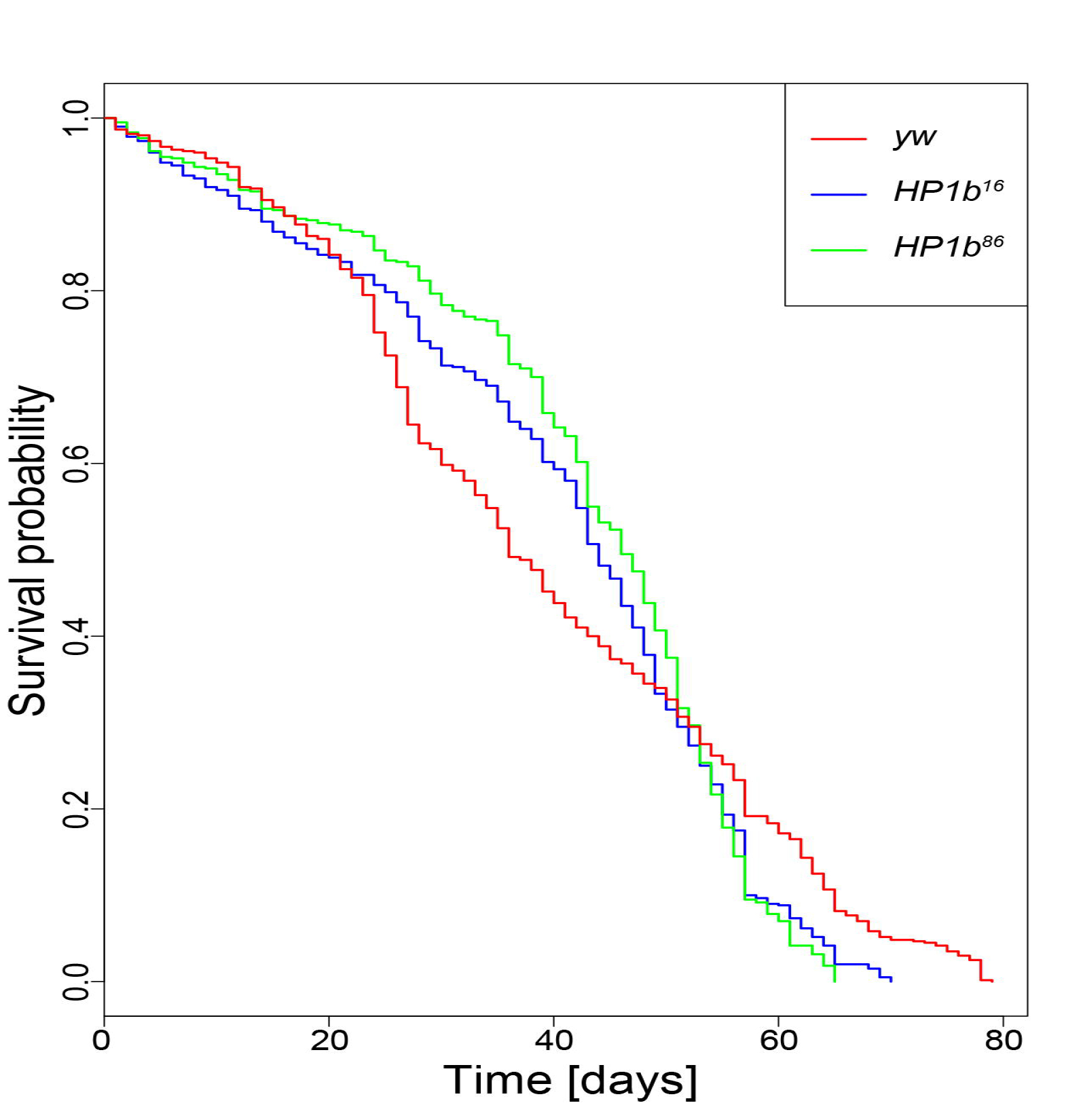
Loss of HP1B impacts survival. Survival curves differ significantly between the *HP1b* mutant strains and the *yw* control strain (p= 0.0483; log rank test). X-axis: Time to death in days. Y-axis: Survival probability. Data shown are combined from three trials, each with 100 animals per genotype/sex. For separate survival curves for males and females, see Figure S4.

### Mutations in *HP1b* lower food intake

As dietary restrictions can result in altered survival curves [58], we investigated if a change in feeding behavior might contribute to the altered survival curves of *HP1b* mutants. We measured food intake using the CAFE assay [39], where the animals are fed a nutrient solution through glass capillaries, recording consumption over an 8hr time period. Representative results from one of three trials are shown in Figure 7A. Both genotype and sex affected the animals’ food intake strongly in all trials (p<0.05, ANOVA). *HP1b* mutant animals consume less food than their *yw* counterparts (Figure 7A), a finding consistent among all three trials despite the fact that individual comparisons are not always significant (Table S1). Across all trials, *HP1b* mutants consumed approximately 30% less food than the *yw* control flies (24-32% depending on trial), and this difference between the *HP1b* mutants and the *yw* control is highly significant in a combined analysis of all three trials (p=0.0000305 for *HP1b^16^* and p=0.0013900 for *HP1b^86^*). The results from this assay indicate that loss of *HP1b* impacts food consumption, suggesting that the physiology and behavior of these animals might have been altered.

**Figure 7.**
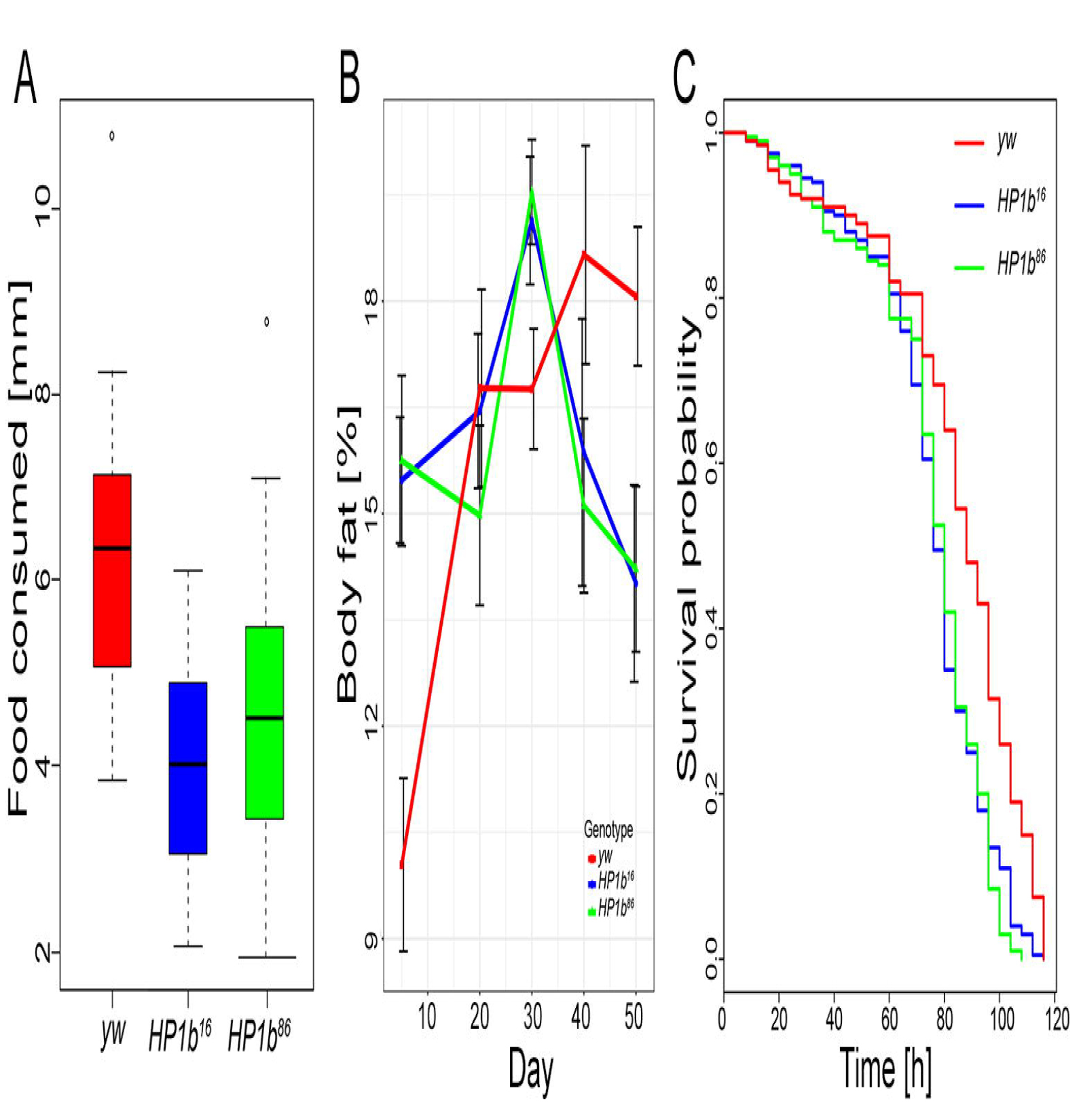
Loss of HP1B reduces food intake, alters body fat levels, and leads to oxidative stress resistance due to altered feeding behavior. **A.** Food intake measured using a CAFÉ assay (Y-axis, decrease in food levels over an 8hr time period as measured by drop in capillary meniscus in mm) reveals lower food intake in *HP1b* mutants (*HP1b^16^*: p=0.0003155; *HP1b^86^*: p=0.0095041; Tukey’s multiple comparisons of means; n=8 per sex and genotype). Data from one representative trial, both sexes combined are shown. For results from the additional trials and separated by sex, see Table S1. **B.** Percent body fat (Y-axis) was measured by QMR in animals of different ages (X-axis). Body fat levels are increased in mutant animals early in life but decrease at a rapid rate after midlife (30-40 days). Red: *yw*; blue: *HP1b^16^*; green: *HP1b^86^*. Error bars: SD. N=4 per sex and genotype. **C.** After paraquat concentration is adjusted to take into account differences in feeding behavior between *HP1b* mutants (blue, green) and the *yw* control strain (red), *HP1b* mutants no longer exhibit increased resistance to oxidative stress. X-axis: Time to death in hours. Y-axis: Survival probability. For separate survival curves for males and females, see Figure S5.

### *HP1b* mutants have altered body fat level

The lower activity levels observed in *HP1b* mutants together with the change in feeding behavior suggests that *HP1b* mutants might have altered body composition as well. We used quantitative magnetic resonance (QMR) to estimate fat and lean body mass. 3-5 day-old *HP1b* mutant flies showed significantly increased fat percentage compared to *yw* (15.5% vs 15.7% versus 10.0%; *HP1b^16^* and *yw* p=0.003, *HP1b^86^* and *yw* p=0.002; Tukey’s multiple comparisons of means; Figure 7B). This increase in body fat content is observed in both male and female animals, but in males, only the *HP1b^16^/yw* comparison reaches statistical significance.

Next, we extended this assay to 50 days, measuring body fat content at 20, 30, 40, and 50 days of age. Specifically, we were interested in determining if the body fat levels remained elevated in the *HP1b* mutant animals, or if a loss of body fat might be responsible for the steep decline in survival seen for these animals around day 40 (Figure 6). Interestingly, the body fat content of *yw* animals had a trajectory different from animals lacking HP1B. In *yw* animals, body fat levels increased with age, from 10.0% at the earliest time point to approximately 18% at 40 and 50 days of age. In contrast, the body fat content of the *HP1b* mutant animals peaks at day 30 at 19% and drops to 14% at day 50. Thus, the level of body fat at day 50 is significantly lower in *HP1b* mutant animals than in the *yw* control strain (*HP1b^16^*: p=0.0111651; *HP1b^86^*: p=0.0159584; Tukey’s multiple comparisons of means). Together, these results demonstrate that loss of HP1B impacts body composition, shifting overall composition towards increased fat levels early in life and experiencing a sharper drop in fat levels during mid-life.

### Oxidative stress resistance in *HP1b* mutants is due to altered feeding behavior

Given that our assays indicate that *HP1b* mutant animals have altered body fat content and altered behavior, both in terms of activity levels and in terms of food intake, we wondered if these changes might explain the increased stress resistance we had observed. Having a higher body fat content and lower energy expenditures due to decreased activity levels could explain the increased starvation resistance of *HP1b* mutants (Figure 5A), while the lower food intake could result in an overall lower dose of paraquat compared to *yw*, thus allowing the *HP1b* mutants to survive longer under oxidative stress (Figure 5B). We tested this hypothesis by carrying out a second oxidative stress assay, this time adjusting the paraquat concentration in the food to account for the reduced feeding rate in *HP1b* mutants. Compared to the initial assay (Figure 5A), the survival curve of *yw* shifts to the right, with *yw* animals now living slightly longer than the *HP1b* mutant animals (Figure 7C). Animals from the *yw* line die on average 83.74h after the initiation of oxidative stress, while *HP1b^16^* animals die on average after 75.08h and *HP1b^86^* after 74.52h (*HP1b^16^*: p= 9.693e-07; *HP1b^86^*: p= 6.548e-07; Kruskal-Wallis rank sum test; Figure 7C). This experiment suggests that the oxidative stress resistance observed for animals lacking HP1B in our initial experiment is due to the decreased food intake level of these animals.

### Expression analysis reveals misregulation of metabolic genes

To gain insights into the causes of the physiological and behavioral changes observed in the animals lacking HP1B, we performed an RNA-seq experiment to compare gene expression levels between the *HP1b* mutants and the *yw* control strain. Using a significance cut-off of p<0.05 (FDR, Benjamini-Hochberg), we find 2179 genes in *HP1b^16^* and 934 genes in *HP1b^86^* that differ significantly from *yw* in their expression levels (Figure S6A). Of these differentially expressed genes, 763 are significantly altered in both *HP1b* mutant strains (Figure S6A). Overall, if we consider all genes, gene expression changes observed between *yw* and *HP1b* mutants occur in roughly equal proportions in either direction (*HP1b^16^*: 46% up, 54% down; *HP1b^86^*: 56% up, 44% down; Figure S6B). In contrast, among the genes with significantly altered gene expression, upregulation is more common (Figure S6B). For example, in the gene set with significantly altered gene expression in both *HP1b* mutant strains, 85% of genes are upregulated, and only 15% are downregulated (Figure S6B). Together, these data illustrate that loss of HP1B impacts a large number of genes and that while upregulation appears more common, gene expression changes can occur in both directions.

Next, we investigated the types of genes affected by loss of HP1B. By analyzing gene ontology (GO) terms associated with the differentially expressed genes using PANTHER [41], we identified a variety of GO terms enriched in the gene set affected by the loss of HP1B (Figure 8). While none of the GO SLIM terms were enriched significantly in the downregulated gene set, the upregulated genes were highly enriched for a variety of metabolism-related terms. These terms include the broad term “metabolic process”, but also more specific terms such as “carbohydrate metabolic process”, “lipid metabolic process”, and “steroid metabolic process” (Figure 8, Table S2). This preponderance of metabolism-related terms is also evident in the full GO analysis (Tables S3 and S4). Thus, the gene expression analysis suggests that loss of HP1B leads to a profoundly altered metabolism which might be the root of the various other phenotypes of *HP1b* mutants noted above.

**Figure 8.**
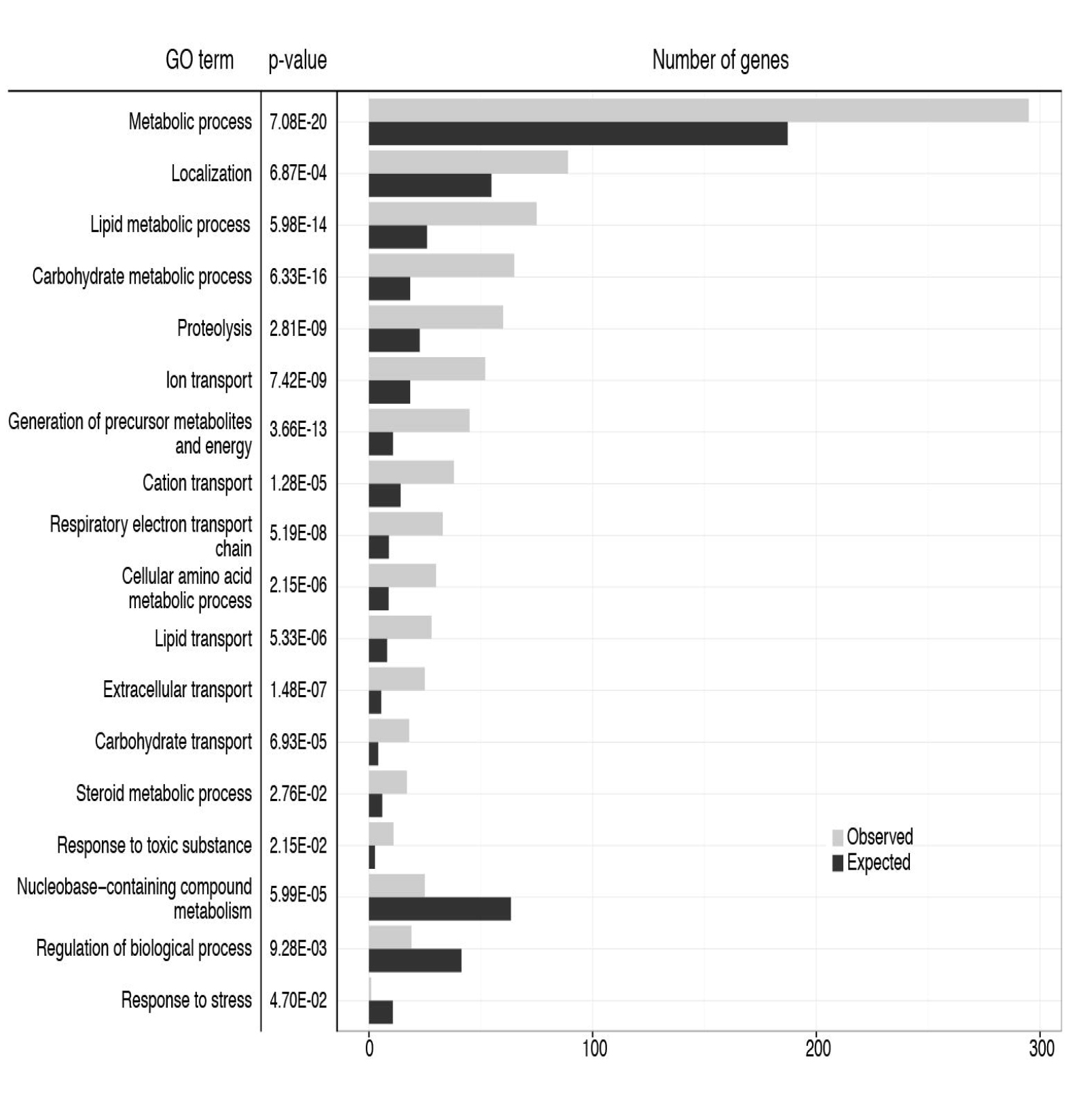
Expression analysis identifies metabolism genes as altered in *HP1b* mutants. GO terms significantly enriched (top) or depleted (bottom three terms) are reported along with the relevant p-values. The bar graphs show the observed (grey) and expected (black) number of genes associated with each GO term. GO terms show are top level terms from GO SLIM. For the results for all GO SLIM terms, see supplemental table X. For the results from the basic GO term analysis see supplemental tables Y (upregulated genes) and Z (downregulated genes).

### Levels of some Krebs cycle metabolites and mitochondrial complexes are lowered in *HP1b* mutants

Given that the expression analysis suggested that metabolic processes were strongly impacted by loss of HP1B, we carried out a targeted metabolite assay for Krebs cycle intermediates using mass spectrometry. Metabolites assayed included pyruvate, lactate, citrate, *cis*-aconitate, isocitrate, 2-hydroxyglutarate, α-keto-gluterate, succinate, fumarate, malate, oxalo-acetate, glutamate, glutamine, and aspartate. Of these compounds, *cis*-aconitate, isocitrate, α-keto-gluterate, and oxalo-acetate were below the limit of quantification. Among the remaining metabolites, citrate and malate levels are impacted by loss of HP1B (Figure 9). Citrate levels in extracts from *HP1b* mutant animals are lower than the citrate levels in the *yw* control samples (p<0.05, t-test, Figure 9A). Malate levels are also lower in the extracts from animals lacking HP1B, but the difference is not significant (p=0.067159718, t-test, Figure 9B), most likely due to the relatively small sample size. Together, these results suggest that the levels of some Krebs cycle metabolites are indeed impacted by the loss of HP1B.

**Figure 9.**
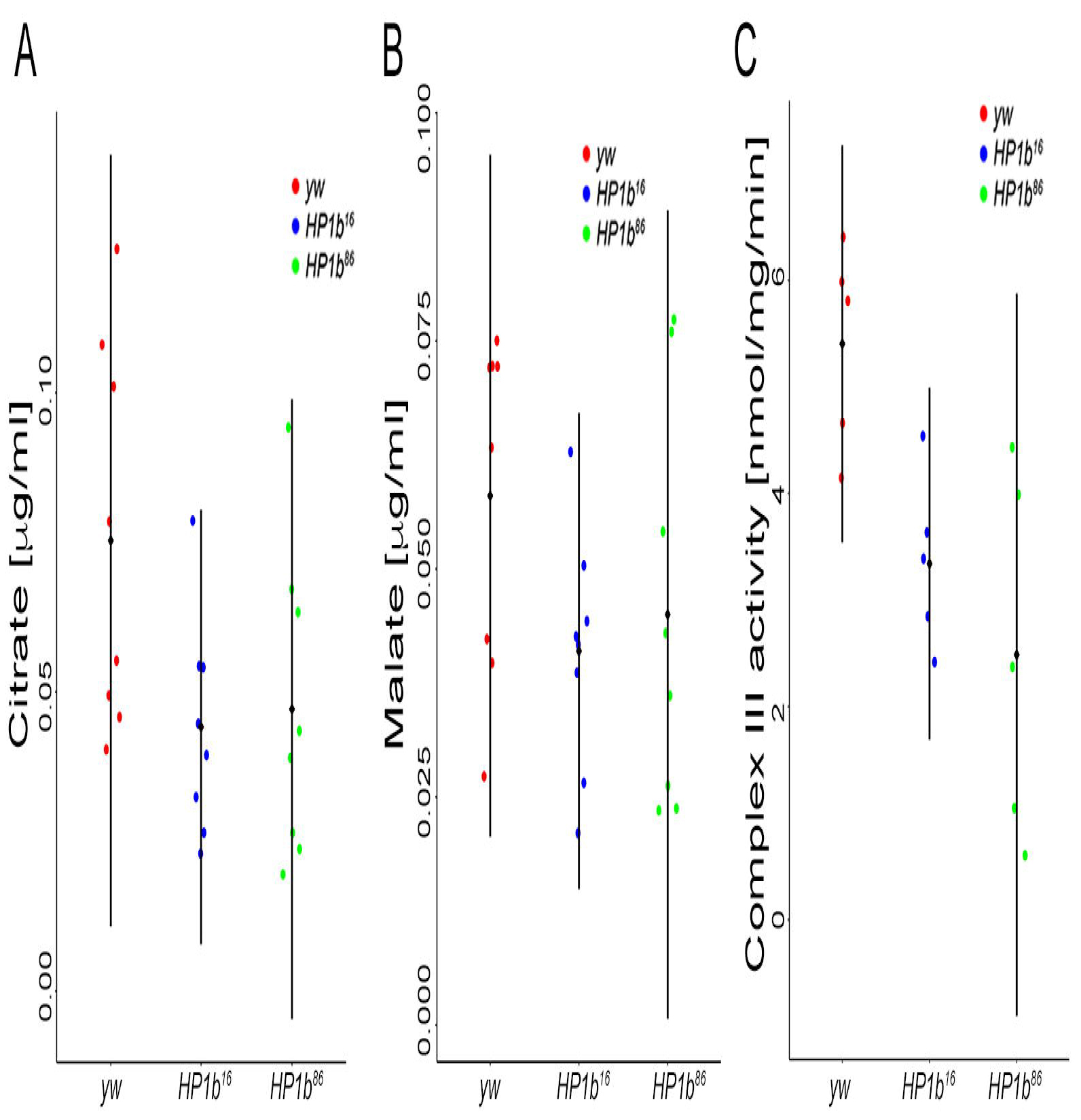
Krebs cycle metabolites and mitochondrial complex III activity are impacted by loss of HP1B. **A&B.** Assay of Krebs cycle metabolites by mass spectroscopy reveals that citrate (**A**) and malate (**B**) levels are lower in *HP1b* mutants. X-axis: Genotypes. Y-axis: Metabolite levels in μg/ml. n=4 per sex and genotype. **C.** Mitochondrial function assays indicate that Complex III activity is reduced in *HP1b* mutant animals (males). X-axis: Genotypes. Y-axis: enzyme activity in mmol/mg/min. n=5 per genotype.

Because both our metabolite and RNA-seq analysis indicated that the mitochondria might be impacted by loss of HP1B, we next assayed two mitochondrial enzymes for their activity, citrate synthase (chosen based on the results from the metabolite analysis) and mitochondrial complex III, which has been linked to an inability to exercise in humans [59]. We found that, while citrate synthase activity levels were similar for HP1B mutant and yw control animals (ANOVA, p>0.05), *HP1b* mutants had decreased levels of complex III activity (Figure 9C, ANOVA, *HP1b^16^* and *yw* p=0.048, *HP1b^86^* and *yw:* p =0.006). On average, the activity measured for complex III in *yw* was 5.4nmol/mg/min (+/- 0.9), while the average for *HP1b* mutants was 3.3nmol/mg/min (+/-0.8, *HP1b^16^*) and 2.5nmol/mg/min (+/-1.7; *HP1b^86^*). Thus, complex III activity was reduced by approximately 48% in the mutant animals compared to the controls. These results demonstrate that loss of HP1B significantly impacts mitochondrial activity, which might explain the lower activity levels of the animals.

## DISCUSSION

In our study, we describe for the first time the phenotypes associated with null mutations in *HP1b*, a member of the HP1 family in Drosophila. Our results indicate that HP1B is not essential for survival, but nonetheless impacts important organism-level phenotypes. While we find that females lacking HP1B produce fewer adult offspring than control animals, homozygous *HP1b* mutants can be maintained as a healthy stock. These findings are consistent with the results of an RNAi knockdown study carried out in 2011, which found slightly decreased viability in flies with lower levels of HP1B [60]. With respect to viability, HP1B thus is similar to HP1C, another somatic HP1 homolog in Drosophila. While *HP1c* null alleles typically are maintained in a heterozygous stock, homozygous mutant flies survive to adulthood and are fertile [61]. The non-essential nature of HP1B and HP1C contrasts with the lethality seen for HP1a, which mainly functions in heterochromatin [24, 25, 48]. Thus, these observations indicate that the phenotypes of null mutations in the Drosophila HP1 proteins reflect their functional divergence.

While not significantly impacting stock viability, phenotypic characterization of *HP1b* mutants revealed that loss of this chromatin protein has far-reaching consequences. The animals exhibit changed behaviors such as lowered food intake and lowered activity levels throughout development. These behavioral alterations lead to a shift in body composition, with the animals accumulating high body fat levels especially early in life. Together, these changes result in two stress resistance phenotypes, starvation stress resistance and oxidative stress resistance. Contrary to common belief, these stress resistance phenotypes are not due to an up-regulated stress response pathway; they are due to altered behavior and body composition: lower activity levels and higher initial body fat content allows the *HP1b* mutant animals to survive longer under starvation conditions, while the lower food intake allows them to better tolerate the oxidative stress conditions as they ingest less oxidizer.

Currently, it is unclear how often phenotypes like the ones described here for *HP1b* mutants occur in other chromatin mutants as few studies go beyond characterizing viability and fertility/offspring number. Fertility defects are seen in a number of chromatin mutants, including *egg* (the Drosophila SETDB1 homolog; [62, 63]), *su(Hw)* (an insulator protein; [64, 65]) and *rhino* (a female germline specific HP1 homolog; [66]). In mutants lacking the H3K9 histone methyltransferase SU(VAR)3-9 or the H3S10 kinase JIL-1, fewer than the expected offspring number are seen, suggesting that fertility and viability are affected [67, 68]. Given the frequency in which fertility and survival are affected by mutations in chromatin components, it is likely that to date unreported phenotypes similar to those found in *HP1b* mutants might occur as well.

The RNA-seq analysis suggests that the various phenotypes of *HP1b* mutants originate in their altered metabolism. This hypothesis is supported by the large fraction of genes associated with metabolism GO terms that are found to be misregulated in the RNA-seq study, the results of the targeted metabolite study, and the mitochondria function assay. To date, no other metabolism studies have been reported for mutants lacking HP1 proteins. However, an RNAi knockdown study of HP1 proteins in Drosophila tissue culture cells found a significant enrichment of genes associated with metabolism (based on GO terms) among the misregulated gene set [69]. These results indicate that the link between HP1B and metabolism is conserved in the tissue culture system as well as animals.

Results from other model organisms further support a link between HP1 proteins and metabolism. Mice, as humans, have three HP1 homologs, HP1α, HP1β, and HP1γ (CBX5, CBX1, CBX3). A hypomorphic allele of the HP1 homolog *Cbx3* (HP1γ) caused death in most mice prior to weaning and is associated with small pup size and altered energy homeostasis [70]. In *C. elegans*, loss of the HP1 homolog HP1L-2 leads to altered lipid metabolism in the animals [71]. These examples illustrate that the link identified in this study between an HP1 protein and metabolism might exist in other species as well.

The mechanism by which HP1B exerts its influence on metabolism genes is currently unclear. The finding that HP1B acts as E(var) in this study would suggest that it has a transcription activating function. However, all reporter genes tested in this study were located in heterochromatin, and when a *lacI* tagged version of HP1B was targeted to two euchromatic transgenes, silencing was reported [52]. Gene expression studies presented here as well as previous expression microarray [69] and qPCR studies [60] also show varying results in that loss of HP1B can lead to increased as well as decreased gene expression. Together, these findings suggest that the impact of HP1B loss on gene expression might be context-dependent.

Context-dependent impacts on gene expression have also been reported for another HP1 protein, HP1a in Drosophila. HP1a is mostly a transcriptional repressor [72], but it also acts as transcriptional activator for approximately 100 euchromatic loci [73] as well as several heterochromatic loci [74-77]. Similar complex associations between HP1 proteins and transcription have been described in other species including mammals [reviewed in [78]], indicating that context-dependent roles in gene regulation are a common characteristic of HP1 proteins.

The complex impacts of HP1 proteins on gene expression might be explained in part by their diverse set of interacting proteins. Interactors include activators, repressors, and RNA pol II itself among many others, including other HP1 proteins [for data from Drosophila, see [79, 80]; reviewed in [78]]. Currently, it is mostly unclear what happens to protein partners if individual HP1 proteins are depleted, and how this depletion affects the various protein complexes HP1 proteins participate in. Thus, knowledge of what additional protein interactions occur at individual HP1 protein binding sites might be necessary to predict the effects of HP1 proteins – and that of their removal - on gene expression at the target site. Further investigation of these interactors might also reveal how HP1B - and other HP proteins - are recruited to their target sites and how their regulation differs among HP1 paralogs. These studies will be essential to determine how the chromatin protein HP1B is linked to the control of metabolism as demonstrated in our studies.

## ACKNOWLEDGEMENTS

We would like to acknowledge SCR Elgin for her support of the initial stages of this project. We also thank the members of the Riddle lab for assistance with fly maintenance, especially LP Watanabe and J Lammons, and for helpful discussion and comments on the manuscript.

We thank the following UAB core facilities: Bio-Analytical Redox Biology core (supported by grant number P30 DK079626/DK/NIDDK from the NIH) for their assistance with the mitochondria assays; Targeted Metabolomics and Proteomics Laboratory core, specifically Taylor Berryhill, (supported by the UAB Health Services Foundation General Endowment Fund Grant and the UAB O’Brien Acute Kidney Injury Center NIH P30 DK079337) for their assistance with the metabolite analysis; UAB Small Animal Phenotyping Core (supported by the NIH Nutrition & Obesity Research Center NIH P30DK056336, UAB Diabetes Research Center NIH P30DK079626, and Nathan Shock Center NIH PAG050886A) for their assistance with the body composition analysis; and Heflin Center for Genomic Science Genomics Core Laboratory (supported by the UAB Comprehensive Cancer Center NIH NCI P30 CA13148 and the UAB Center for AIDS Research NIH P30 AI027767) for their assistance with the RNA-seq experiments.

Stocks obtained from the Bloomington Drosophila Stock Center (NIH P40OD018537) were used in this study.

The JLA20 monoclonal antibody developed by J. J.-C. Lin was obtained from the Developmental Studies Hybridoma Bank, created by the NICHD of the NIH and maintained at The University of Iowa, Department of Biology, Iowa City, IA 52242.

This material is based upon work supported by the National Science Foundation under Grant No. 1552586 (NCR). Additional funding was provided by American Cancer Society Institutional Research Grant to NCR.

## SUPPORTING INFORMATION

**Table S1. Genotype and sex strongly impact feeding behavior.**

**Table S2. Expanded GO analysis results I.**

**Table S3. Expanded GO analysis results II.**

**Table S4. Expanded GO analysis results III.**

**Figure S1. *HP1b* mutations enhance *Sb^v^* variegation, but can act as Su(var) or E(var), depending on the reporter insertion site of a *white* reporter. A.** Eye pigment assays for females from the five reporter insertions assayed. The response of an *hsp70-white* reporter to a single mutant *HP1b* allele in female flies depends on the reporter location. *: comparisons of mutant eye pigment levels to the *y^-^w^-^* control that are statistically significant (p<0.05, t-test). Y-axis: OD_480_ measuring eye pigment. X-axis: reporter insertions. Results from the *y^-^w^-^* control are in red, from *HP1b^16^* in blue, and from *HP1b^86^* in green. Box plots: black bar - median; box - +25% and −25% quartiles; whiskers – maximum and minimum; circles – outliers; n=6-10. **B.** *Stubble* variegation assays indicate that *HP1b* is an enhancer of variegation, as the presence of the *HP1b* mutant alleles significantly increases the number of wildtype bristles compared to the *y^-^w^-^* control (p<0.01, Tukey multiple comparisons of means; n=19-39). Data from females. Results from the *y^-^w^-^* control are in red, from *HP1b^16^* in blue, and from *HP1b^86^* in green. Box plots: black bar - median; box - +25% and −25% quartiles; whiskers – maximum and minimum; circles – outliers.

**Figure S2. Adult activity levels are impacted in both males and females. A.** Female flies lacking HP1B show significantly lower activity levels than animals of the *yw* control genotype (p<0.001 for both comparisons; Tukey’s HSD). **B.** In males, the activity levels of animals lacking HP1B is reduced compared to *yw*, but the effect is less pronounced than in females and only significant for the *HP1b^86^* allele (p-value not significant for *HP1b^16^*, and p<0.001 for *HP1b^86^*, Tukey’s HSD). Y-axis: activity level as measured by recorded beam crossings in an activity monitor for 10 flies within the 2hr assay period. Black diamond and bar: mean +/- SD. N=25 per sex/genotype combination.

**Figure S3. Loss of HP1B impacts survival during stress in both males and females. A.** Female *HP1b* mutant animals survive significantly longer during starvation condition than *yw* control animals (top; p = 6.455e-09 and p = 4.672e-07 for *HP1b^16^* and *HP1b^86^* respectively; Kruskal-Wallis rank sum test), while for males, only the comparison between *HP1b^16^* and *yw* is significant (bottom; p = 2.054e-07; Kruskal-Wallis rank sum test). **B.** Female *HP1b* mutant animals survive significantly longer after exposure to the oxidizer paraquat than *yw* control animals (top; p = 3.752e-10 and p < 2.2e-16 for *HP1b^16^* and *HP1b^86^* respectively; Kruskal-Wallis rank sum test), as do males (bottom; p =1.252e-05 and p = 1.79e-14 for *HP1b^16^* and *HP1b^86^* respectively; Kruskal-Wallis rank sum test). **C.** *HP1b* mutant animals, both male and female, do not show improved survival during heat stress conditions (37°C) compared to *yw* control animals. In contrast, *HP1b^86^* shows significantly lower survival (p = 0.001803 in females [top] and p = 1.499e-08in males [bottom]; Kruskal-Wallis rank sum test).

For **A-C**: X-axis - time to death in hours. Y-axis - survival probability. Data shown are from three trials, each with 100 animals per genotype.

**Figure S4. Loss of HP1B impacts the survival curve of females and males differently. A.** Survival curves differ significantly between the *HP1b* mutant strains and the *yw* control strain in females (*HP1b^16^*: p= 0.0259; *HP1b^86^*: p= 0.00563; log rank test). **B.** In males, the survival curves of *HP1b* mutant animals do not differ significantly from the survival curves of *yw* control animals (p not significant; log rank test).

**A+B**: X-axis - time to death in days. Y-axis - survival probability. Data shown are combined from three trials, each with 100 animals per genotype/sex.

**Figure S5. Altered feeding behavior is the likely cause of increased oxidative stress resistance as measured by paraquat feeding in both males and females.** The paraquat concentration was adjusted to take into account differences in feeding behavior between *HP1b* mutants (blue, green) and the *yw* control strain (red). **A. Females.** After adjusting the paraquat concentration, female *HP1b* mutant animals die earlier than their *yw* counterparts (*HP1b^16^*: p= 7.413e-11; *HP1b^86^*: p= 2.113e-14; Kruskal-Wallis rank sum test). **B. Males.** After adjusting the paraquat concentration, no difference in survival is detected between *HP1b* mutants and *yw* (*HP1b^16^*: p= 0.31; *HP1b^86^*: p= 0.9883; Kruskal-Wallis-rank sum test). Y-axis: survival probability; X-axis: Time to death in hours. N=100 animals per genotype/sex.

**Figure S6. Gene expression studies identify a large number of genes impacted by loss of HP1B. A.** Venn diagram illustrating the overlap in the gene sets identified as significantly altered in gene expression in the *HP1b^16^* and *HP1b^86^* mutant strains. Only genes with an FDR<0.05 are included (Benjamini-Hochberg). **B.** Stacked bar graph illustrating the percentage of genes up-versus down-regulated upon loss of HP1B compared to the *yw* control strain. Red: upregulated genes; blue: downregulated genes.

## REFERENCES

1. Filion GJ, van Bemmel JG, Braunschweig U, Talhout W, Kind J, Ward LD, et al. Systematic protein location mapping reveals five principal chromatin types in Drosophila cells. Cell. 2010;143(2):212–24. doi: 10.1016/j.cell.2010.09.009. PubMed PMID: 20888037; PubMed Central PMCID: PMC3119929.

2. Heintzman ND, Stuart RK, Hon G, Fu Y, Ching CW, Hawkins RD, et al. Distinct and predictive chromatin signatures of transcriptional promoters and enhancers in the human genome. Nature genetics. 2007;39(3):311–8. doi: 10.1038/ng1966. PubMed PMID: 17277777.

3. Kharchenko PV, Alekseyenko AA, Schwartz YB, Minoda A, Riddle NC, Ernst J, et al. Comprehensive analysis of the chromatin landscape in Drosophila melanogaster. Nature. 2011;471(7339):480–5. Epub 2010/12/24. doi: nature09725 [pii] 10.1038/nature09725. PubMed PMID: 21179089; PubMed Central PMCID: PMC3109908.

4. Grewal SI, Elgin SC. Transcription and RNA interference in the formation of heterochromatin. Nature. 2007;447(7143):399–406. doi: 10.1038/nature05914. PubMed PMID: 17522672; PubMed Central PMCID: PMC2950806.

5. Kwon SH, Workman JL. The changing faces of HP1: From heterochromatin formation and gene silencing to euchromatic gene expression: HP1 acts as a positive regulator of transcription. Bioessays. 2011;33(4):280–9. Epub 2011/01/29. doi: 10.1002/bies.201000138. PubMed PMID: 21271610.

6. Vermaak D, Malik HS. Multiple roles for heterochromatin protein 1 genes in Drosophila. Annual review of genetics. 2009;43:467–92. doi: 10.1146/annurev-genet-102108-134802. PubMed PMID: 19919324.

7. Dialynas GK, Vitalini MW, Wallrath LL. Linking Heterochromatin Protein 1 (HP1) to cancer progression. Mutation research. 2008;647(1-2):13–20. doi: 10.1016/j.mrfmmm.2008.09.007. PubMed PMID: 18926834; PubMed Central PMCID: PMC2637788.

8. Wood JG, Hillenmeyer S, Lawrence C, Chang C, Hosier S, Lightfoot W, et al. Chromatin remodeling in the aging genome of Drosophila. Aging cell. 2010;9(6):971–8. doi: 10.1111/j.1474-9726.2010.00624.x. PubMed PMID: 20961390; PubMed Central PMCID: PMC2980570.

9. Zhang R, Adams PD. Heterochromatin and its relationship to cell senescence and cancer therapy. Cell cycle. 2007;6(7):784–9. PubMed PMID: 17377503.

10. Gaudin V, Libault M, Pouteau S, Juul T, Zhao G, Lefebvre D, et al. Mutations in LIKE HETEROCHROMATIN PROTEIN 1 affect flowering time and plant architecture in Arabidopsis. Development. 2001;128(23):4847–58. PubMed PMID: 11731464.

11. Lomberk G, Wallrath L, Urrutia R. The Heterochromatin Protein 1 family. Genome biology. 2006;7(7):228. doi: 10.1186/gb-2006-7-7-228. PubMed PMID: 17224041; PubMed Central PMCID: PMC1779566.

12. Aasland R, Stewart AF. The chromo shadow domain, a second chromo domain in heterochromatin-binding protein 1, HP1. Nucleic acids research. 1995;23(16):3168–73. PubMed PMID: 7667093; PubMed Central PMCID: PMC307174.

13. Bannister AJ, Zegerman P, Partridge JF, Miska EA, Thomas JO, Allshire RC, et al. Selective recognition of methylated lysine 9 on histone H3 by the HP1 chromo domain. Nature. 2001;410(6824):120–4. doi: 10.1038/35065138. PubMed PMID: 11242054.

14. Jacobs SA, Khorasanizadeh S. Structure of HP1 chromodomain bound to a lysine 9-methylated histone H3 tail. Science. 2002;295(5562):2080–3. doi: 10.1126/science.1069473. PubMed PMID: 11859155.

15. Jacobs SA, Taverna SD, Zhang Y, Briggs SD, Li J, Eissenberg JC, et al. Specificity of the HP1 chromo domain for the methylated N-terminus of histone H3. Embo J. 2001;20(18):5232–41. PubMed PMID: 11566886.

16. Lachner M, O’Carroll D, Rea S, Mechtler K, Jenuwein T. Methylation of histone H3 lysine 9 creates a binding site for HP1 proteins. Nature. 2001;410(6824):116–20. doi: 10.1038/35065132. PubMed PMID: 11242053.

17. Nielsen PR, Nietlispach D, Mott HR, Callaghan J, Bannister A, Kouzarides T, et al. Structure of the HP1 chromodomain bound to histone H3 methylated at lysine 9. Nature. 2002;416(6876):103–7. doi: 10.1038/nature722. PubMed PMID: 11882902.

18. Shanle EK, Shinsky SA, Bridgers JB, Bae N, Sagum C, Krajewski K, et al. Histone peptide microarray screen of chromo and Tudor domains defines new histone lysine methylation interactions. Epigenetics Chromatin. 2017;10:12. doi: 10.1186/s13072-017-0117-5. PubMed PMID: 28293301; PubMed Central PMCID: PMCPMC5348760.

19. Smothers JF, Henikoff S. The hinge and chromo shadow domain impart distinct targeting of HP1-like proteins. Molecular and cellular biology. 2001;21(7):2555–69. doi: 10.1128/MCB.21.7.2555-2569.2001. PubMed PMID: 11259603; PubMed Central PMCID: PMC86887.

20. Brasher SV, Smith BO, Fogh RH, Nietlispach D, Thiru A, Nielsen PR, et al. The structure of mouse HP1 suggests a unique mode of single peptide recognition by the shadow chromo domain dimer. The EMBO journal. 2000;19(7):1587–97. doi: 10.1093/emboj/19.7.1587. PubMed PMID: 10747027; PubMed Central PMCID: PMC310228.

21. Cowieson NP, Partridge JF, Allshire RC, McLaughlin PJ. Dimerisation of a chromo shadow domain and distinctions from the chromodomain as revealed by structural analysis. Current biology: CB. 2000;10(9):517–25. PubMed PMID: 10801440.

22. Nielsen AL, Oulad-Abdelghani M, Ortiz JA, Remboutsika E, Chambon P, Losson R. Heterochromatin formation in mammalian cells: interaction between histones and HP1 proteins. Molecular cell. 2001;7(4):729–39. PubMed PMID: 11336697.

23. Thiru A, Nietlispach D, Mott HR, Okuwaki M, Lyon D, Nielsen PR, et al. Structural basis of HP1/PXVXL motif peptide interactions and HP1 localisation to heterochromatin. The EMBO journal. 2004;23(3):489–99. doi: 10.1038/sj.emboj.7600088. PubMed PMID: 14765118; PubMed Central PMCID: PMC1271814.

24. Eissenberg JC, Morris GD, Reuter G, Hartnett T. The heterochromatin-associated protein HP-1 is an essential protein in Drosophila with dosage-dependent effects on position-effect variegation. Genetics. 1992;131(2):345–52. PubMed PMID: 1644277; PubMed Central PMCID: PMC1205009.

25. James TC, Eissenberg JC, Craig C, Dietrich V, Hobson A, Elgin SC. Distribution patterns of HP1, a heterochromatin-associated nonhistone chromosomal protein of Drosophila. Eur J Cell Biol. 1989;50(1):170–80. PubMed PMID: 2515059.

26. Vermaak D, Henikoff S, Malik HS. Positive selection drives the evolution of rhino, a member of the heterochromatin protein 1 family in Drosophila. PLoS genetics. 2005;1(1):96–108. doi: 10.1371/journal.pgen.0010009. PubMed PMID: 16103923; PubMed Central PMCID: PMC1183528.

27. Levine MT, McCoy C, Vermaak D, Lee YC, Hiatt MA, Matsen FA, et al. Phylogenomic analysis reveals dynamic evolutionary history of the Drosophila heterochromatin protein 1 (HP1) gene family. PLoS genetics. 2012;8(6):e1002729. doi: 10.1371/journal.pgen.1002729. PubMed PMID: 22737079; PubMed Central PMCID: PMC3380853.

28. Minc E, Allory Y, Worman HJ, Courvalin JC, Buendia B. Localization and phosphorylation of HP1 proteins during the cell cycle in mammalian cells. Chromosoma. 1999;108(4):220–34. PubMed PMID: 10460410.

29. Shaffer CD, Wuller JM, Elgin SC. Raising large quantities of Drosophila for biochemical experiments. Methods in cell biology. 1994;44:99–108. PubMed PMID: 7707979.

30. Buszczak M, Paterno S, Lighthouse D, Bachman J, Planck J, Owen S, et al. The carnegie protein trap library: a versatile tool for Drosophila developmental studies. Genetics. 2007;175(3):1505–31. doi: 10.1534/genetics.106.065961. PubMed PMID: 17194782; PubMed Central PMCID: PMC1840051.

31. Team RC. R: A Language and Environment for Statistical Computing. R Foundation for Statistical Computing; 2015.

32. Khesin RB, Leibovitch BA. Influence of deficiency of the histone gene-containing 38B-40 region on X-chromosome template activity and the white gene position effect variegation in Drosophila melanogaster. Molecular & general genetics: MGG. 1978;162(3):323–8. PubMed PMID: 98701.

33. Csink AK, Linsk R, Birchler JA. The Lighten up (Lip) gene of Drosophila melanogaster, a modifier of retroelement expression, position effect variegation and white locus insertion alleles. Genetics. 1994;138(1):153–63. PubMed PMID: 8001783; PubMed Central PMCID: PMCPMC1206127.

34. Tettweiler G, Miron M, Jenkins M, Sonenberg N, Lasko PF. Starvation and oxidative stress resistance in Drosophila are mediated through the eIF4E-binding protein, d4E-BP. Genes & development. 2005;19(16):1840–3. doi: 10.1101/gad.1311805. PubMed PMID: 16055649; PubMed Central PMCID: PMCPMC1186182.

35. Therneau T. A package for survival analysis in S. https://CRAN.R-project.org/package=survival2015.

36. Wang HD, Kazemi-Esfarjani P, Benzer S. Multiple-stress analysis for isolation of Drosophila longevity genes. Proceedings of the National Academy of Sciences of the United States of America. 2004;101(34):12610–5. doi: 10.1073/pnas.0404648101. PubMed PMID: 15308776; PubMed Central PMCID: PMCPMC515105.

37. Rzezniczak TZ, Douglas LA, Watterson JH, Merritt TJ. Paraquat administration in Drosophila for use in metabolic studies of oxidative stress. Anal Biochem. 2011;419(2):345–7. doi: 10.1016/j.ab.2011.08.023. PubMed PMID: 21910964.

38. Linford NJ, Bilgir C, Ro J, Pletcher SD. Measurement of lifespan in Drosophila melanogaster. J Vis Exp. 2013;(71). doi: 10.3791/50068. PubMed PMID: 23328955; PubMed Central PMCID: PMCPMC3582515.

39. Ja WW, Carvalho GB, Mak EM, de la Rosa NN, Fang AY, Liong JC, et al. Prandiology of Drosophila and the CAFE assay. Proceedings of the National Academy of Sciences of the United States of America. 2007;104(20):8253–6. doi: 10.1073/pnas.0702726104. PubMed PMID: 17494737; PubMed Central PMCID: PMCPMC1899109.

40. Trapnell C, Roberts A, Goff L, Pertea G, Kim D, Kelley DR, et al. Differential gene and transcript expression analysis of RNA-seq experiments with TopHat and Cufflinks. Nat Protoc. 2012;7(3):562–78. doi: 10.1038/nprot.2012.016. PubMed PMID: 22383036; PubMed Central PMCID: PMCPMC3334321.

41. Mi H, Huang X, Muruganujan A, Tang H, Mills C, Kang D, et al. PANTHER version 11: expanded annotation data from Gene Ontology and Reactome pathways, and data analysis tool enhancements. Nucleic acids research. 2017;45(D1):D183–D9. doi: 10.1093/nar/gkw1138. PubMed PMID: 27899595; PubMed Central PMCID: PMCPMC5210595.

42. Xia J, Wishart DS. Using MetaboAnalyst 3.0 for Comprehensive Metabolomics Data Analysis. Curr Protoc Bioinformatics. 2016;55:14 0 1–0 91. doi: 10.1002/cpbi.11. PubMed PMID: 27603023.

43. Miwa S, St-Pierre J, Partridge L, Brand MD. Superoxide and hydrogen peroxide production by Drosophila mitochondria. Free Radic Biol Med. 2003;35(8):938–48. PubMed PMID: 14556858.

44. Lowry OH, Rosebrough NJ, Farr AL, Randall RJ. Protein measurement with the Folin phenol reagent. J Biol Chem. 1951;193(1):265–75. PubMed PMID: 14907713.

45. Leek BT, Mudaliar SR, Henry R, Mathieu-Costello O, Richardson RS. Effect of acute exercise on citrate synthase activity in untrained and trained human skeletal muscle. Am J Physiol Regul Integr Comp Physiol. 2001;280(2):R441–7. PubMed PMID: 11208573.

46. Srere P. Citrate synthase. Methods in enzymology. 1969;13:3–5.

47. Spinazzi M, Casarin A, Pertegato V, Salviati L, Angelini C. Assessment of mitochondrial respiratory chain enzymatic activities on tissues and cultured cells. Nat Protoc. 2012;7(6):1235–46. doi: 10.1038/nprot.2012.058. PubMed PMID: 22653162.

48. Eissenberg JC, James TC, Foster-Hartnett DM, Hartnett T, Ngan V, Elgin SC. Mutation in a heterochromatin-specific chromosomal protein is associated with suppression of position-effect variegation in Drosophila melanogaster. Proceedings of the National Academy of Sciences of the United States of America. 1990;87(24):9923–7. PubMed PMID: 2124708; PubMed Central PMCID: PMC55286.

49. St Pierre SE, Ponting L, Stefancsik R, McQuilton P, FlyBase C. FlyBase 102--advanced approaches to interrogating FlyBase. Nucleic acids research. 2014;42(Database issue):D780–8. doi: 10.1093/nar/gkt1092. PubMed PMID: 24234449; PubMed Central PMCID: PMC3964969.

50. Graveley BR, Brooks AN, Carlson JW, Duff MO, Landolin JM, Yang L, et al. The developmental transcriptome of Drosophila melanogaster. Nature. 2011;471(7339):473–9. doi: 10.1038/nature09715. PubMed PMID: 21179090; PubMed Central PMCID: PMC3075879.

51. mod EC, Roy S, Ernst J, Kharchenko PV, Kheradpour P, Negre N, et al. Identification of functional elements and regulatory circuits by Drosophila modENCODE. Science. 2010;330(6012):1787–97. doi: 10.1126/science.1198374. PubMed PMID: 21177974; PubMed Central PMCID: PMC3192495.

52. Font-Burgada J, Rossell D, Auer H, Azorin F. Drosophila HP1c isoform interacts with the zinc-finger proteins WOC and Relative-of-WOC to regulate gene expression. Genes & development. 2008;22(21):3007–23. Epub 2008/11/05. doi: 10.1101/gad.481408. PubMed PMID: 18981478; PubMed Central PMCID: PMC2577788.

53. Wallrath LL, Elgin SC. Position effect variegation in Drosophila is associated with an altered chromatin structure. Genes Dev. 1995;9(10):1263–77. Epub 1995/05/15. PubMed PMID: 7758950.

54. Clancy DJ, Gems D, Harshman LG, Oldham S, Stocker H, Hafen E, et al. Extension of life-span by loss of CHICO, a Drosophila insulin receptor substrate protein. Science. 2001;292(5514):104–6. doi: 10.1126/science.1057991. PubMed PMID: 11292874.

55. Lin YJ, Seroude L, Benzer S. Extended life-span and stress resistance in the Drosophila mutant methuselah. Science. 1998;282(5390):943–6. PubMed PMID: 9794765.

56. Spencer CC, Howell CE, Wright AR, Promislow DE. Testing an ‘aging gene’ in long-lived drosophila strains: increased longevity depends on sex and genetic background. Aging cell. 2003;2(2):123–30. PubMed PMID: 12882325; PubMed Central PMCID: PMCPMC3991309.

57. Tower J. Heat shock proteins and Drosophila aging. Exp Gerontol. 2011;46(5):355–62. doi: 10.1016/j.exger.2010.09.002. PubMed PMID: 20840862; PubMed Central PMCID: PMCPMC3018744.

58. Partridge L, Piper MD, Mair W. Dietary restriction in Drosophila. Mech Ageing Dev. 2005;126(9):938–50. doi: 10.1016/j.mad.2005.03.023. PubMed PMID: 15935441.

59. DiMauro S, Andreu AL. Mutations in mitochondrial DNA as a cause of exercise intolerance. Ann Med. 2001;33(7):472–6. PubMed PMID: 11680795.

60. Zhang D, Wang D, Sun F. Drosophila melanogaster heterochromatin protein HP1b plays important roles in transcriptional activation and development. Chromosoma. 2011;120(1):97–108. doi: 10.1007/s00412-010-0294-5. PubMed PMID: 20857302.

61. Gramates LS, Marygold SJ, Santos GD, Urbano JM, Antonazzo G, Matthews BB, et al. FlyBase at 25: looking to the future. Nucleic Acids Res. 2017;45(D1):D663–D71. Epub 2016/11/02. doi: 10.1093/nar/gkw1016. PubMed PMID: 27799470; PubMed Central PMCID: PMCPMC5210523.

62. Clough E, Moon W, Wang S, Smith K, Hazelrigg T. Histone methylation is required for oogenesis in Drosophila. Development. 2007;134(1):157–65. PubMed PMID: 17164421.

63. Stabell M, Bjorkmo M, Aalen RB, Lambertsson A. The Drosophila SET domain encoding gene dEset is essential for proper development. Hereditas. 2006;143(2006):177–88. PubMed PMID: 17362353.

64. Baxley RM, Soshnev AA, Koryakov DE, Zhimulev IF, Geyer PK. The role of the Suppressor of Hairy-wing insulator protein in Drosophila oogenesis. Dev Biol. 2011;356(2):398–410. doi: 10.1016/j.ydbio.2011.05.666. PubMed PMID: 21651900; PubMed Central PMCID: PMCPMC3143288.

65. Harrison DA, Gdula DA, Coyne RS, Corces VG. A leucine zipper domain of the suppressor of Hairy-wing protein mediates its repressive effect on enhancer function. Genes & development. 1993;7(10):1966–78. PubMed PMID: 7916729.

66. Volpe AM, Horowitz H, Grafer CM, Jackson SM, Berg CA. Drosophila rhino encodes a female-specific chromo-domain protein that affects chromosome structure and egg polarity. Genetics. 2001;159(3):1117–34. PubMed PMID: 11729157; PubMed Central PMCID: PMCPMC1461866.

67. Peng JC, Karpen GH. H3K9 methylation and RNA interference regulate nucleolar organization and repeated DNA stability. Nature cell biology. 2007;9(1):25–35. PubMed PMID: 17159999.

68. Wang Y, Zhang W, Jin Y, Johansen J, Johansen KM. The JIL-1 tandem kinase mediates histone H3 phosphorylation and is required for maintenance of chromatin structure in Drosophila. Cell. 2001;105(4):433–43. PubMed PMID: 11371341.

69. Lee DH, Li Y, Shin DH, Yi SA, Bang SY, Park EK, et al. DNA microarray profiling of genes differentially regulated by three heterochromatin protein 1 (HP1) homologs in Drosophila. Biochemical and biophysical research communications. 2013;434(4):820–8. doi: 10.1016/j.bbrc.2013.04.020. PubMed PMID: 23611785.

70. Aydin E, Kloos DP, Gay E, Jonker W, Hu L, Bullwinkel J, et al. A hypomorphic Cbx3 allele causes prenatal growth restriction and perinatal energy homeostasis defects. J Biosci. 2015;40(2):325–38. PubMed PMID: 25963260.

71. Meister P, Schott S, Bedet C, Xiao Y, Rohner S, Bodennec S, et al. Caenorhabditis elegans Heterochromatin protein 1 (HPL-2) links developmental plasticity, longevity and lipid metabolism. Genome biology. 2011;12(12):R123. doi: 10.1186/gb-2011-12-12-r123. PubMed PMID: 22185090; PubMed Central PMCID: PMCPMC3334618.

72. Eissenberg JC, Elgin SC. The HP1 protein family: getting a grip on chromatin. Current opinion in genetics & development. 2000;10(2):204–10. PubMed PMID: 10753776.

73. Piacentini L, Fanti L, Negri R, Del Vescovo V, Fatica A, Altieri F, et al. Heterochromatin protein 1 (HP1a) positively regulates euchromatic gene expression through RNA transcript association and interaction with hnRNPs in Drosophila. PLoS genetics. 2009;5(10):e1000670. doi: 10.1371/journal.pgen.1000670. PubMed PMID: 19798443; PubMed Central PMCID: PMC2743825.

74. Cryderman DE, Vitalini MW, Wallrath LL. Heterochromatin protein 1a is required for an open chromatin structure. Transcription. 2011;2(2):95–9. Epub 2011/04/07. doi: 10.4161/trns.2.2.14687. PubMed PMID: 21468237; PubMed Central PMCID: PMCPMC3062402.

75. Eberl DF, Duyf BJ, Hilliker AJ. The role of heterochromatin in the expression of a heterochromatic gene, the rolled locus of Drosophila melanogaster. Genetics. 1993;134(1):277–92. PubMed PMID: 8514136; PubMed Central PMCID: PMC1205430.

76. Howe M, Dimitri P, Berloco M, Wakimoto BT. Cis-effects of heterochromatin on heterochromatic and euchromatic gene activity in Drosophila melanogaster. Genetics. 1995;140(3):1033–45. PubMed PMID: 7672575; PubMed Central PMCID: PMC1206659.

77. Lu BY, Emtage PC, Duyf BJ, Hilliker AJ, Eissenberg JC. Heterochromatin protein 1 is required for the normal expression of two heterochromatin genes in Drosophila. Genetics. 2000;155(2):699–708. PubMed PMID: 10835392; PubMed Central PMCID: PMC1461102.

78. Eissenberg JC, Elgin SC. HP1a: a structural chromosomal protein regulating transcription. Trends Genet. 2014;30(3):103–10. Epub 2014/02/22. doi: 10.1016/j.tig.2014.01.002. PubMed PMID: 24555990; PubMed Central PMCID: PMCPMC3991861.

79. Alekseyenko AA, Gorchakov AA, Zee BM, Fuchs SM, Kharchenko PV, Kuroda MI. Heterochromatin-associated interactions of Drosophila HP1a with dADD1, HIPP1, and repetitive RNAs. Genes & development. 2014;28(13):1445–60. doi: 10.1101/gad.241950.114. PubMed PMID: 24990964; PubMed Central PMCID: PMCPMC4083088.

80. Ryu HW, Lee DH, Florens L, Swanson SK, Washburn MP, Kwon SH. Analysis of the heterochromatin protein 1 (HP1) interactome in Drosophila. Journal of proteomics. 2014;102:137–47. doi: 10.1016/j.jprot.2014.03.016. PubMed PMID: 24681131.

